# MISTERMINATE Mechanistically Links Mitochondrial Dysfunction with Proteostasis Failure

**DOI:** 10.1101/554634

**Authors:** Zhihao Wu, Ishaq Tantray, Junghyun Lim, Songjie Chen, Yu Li, Zoe Davis, Cole Sitron, Jason Dong, Suzana Gispert, Georg Auburger, Onn Brandman, Xiaolin Bi, Michael Snyder, Bingwei Lu

## Abstract

Mitochondrial dysfunction and proteostasis failure frequently coexist as hallmarks of neurodegenerative disease. How these pathologies are related is not well understood. Here we describe a phenomenon termed MISTERMINATE (<u>mi</u>tochondrial <u>s</u>tress-induced translational <u>term</u>ination impairment a<u>n</u>d protein c<u>a</u>rboxyl terminal <u>e</u>xtension), which mechanistically links mitochondrial dysfunction with proteostasis failure. We show that mitochondrial dysfunction impairs translational termination of nuclear-encoded mitochondrial mRNAs including *complex-I 30kD subunit* (*C-I30*) mRNA, occurring on mitochondrial surface in *Drosophila* and mammalian cells. Ribosomes stalled at the normal stop codon continue to add to the C-terminus of C-I30 certain amino acids non-coded by mRNA template. C-terminally-extended C-I30 is toxic when assembled into C-I and forms aggregates in the cytosol. Enhancing co-translational quality control prevents C-I30 C-terminal extension and rescues mitochondrial and neuromuscular degeneration in a Parkinson’s disease model. These findings emphasize the importance of efficient translation termination and reveal unexpected link between mitochondrial health and proteome homeostasis mediated by MISTERMINATE.

## INTRODUCTION

Protein aggregation is a defining feature of age-related neurodegenerative diseases. Hallmark aggregates include those composed of the Aβ peptides in Alzheimer’s disease (AD), α-Syn in Parkinson’s disease (PD), tau in tauopathies, and TDP-43 in amyotrophic lateral sclerosis (ALS) (Hartl, 2017; Chiti and Dobson, 2017). While it remains unclear whether the large aggregates or some smaller intermediates are the toxic species (Hartl, 2017; Sontag et al., 2017), accumulation of faulty proteins seems to play pathogenic roles.

Previous studies of protein aggregation have largely focused on alterations of fully synthesized proteins. Such changes emphasize posttranslational modifications, and significant efforts have been directed toward linking these molecular events with aging-related effects, e.g. oxidative stress and impairments of the autophagy and ubiquitin-proteosome system (UPS) (Bossy-Wetzel et al., 2004; Ciechanover and Brundin, 2003).

Recent studies draw attention to an emerging role of co-translational quality control (QC) in proteome homeostasis (Brandman and Hegde, 2016; Joazeiro, 2017). There is widespread ubiquitination and degradation of nascent peptide chains (NPCs) associated with stalled as well as active translation (Wang et al., 2013a; Duttler et al., 2013). Studies in yeast showed that NPCs on stalled ribosomes can be modified, while still attached to 60S subunits, by C-terminal Ala and Thr addition (CAT-tailing) (Shen et al., 2015). CAT-tailing may enable the degradation of aberrant NPCs by making the most recently added K residues available for ubiquitination (Kostova et al., 2017). While CAT-tailing as part of the ribosome-associated quality control (RQC) complex may induce the heat shock response (Brandman et al., 2012), failure in the timely removal of CAT-tailed proteins can disrupt proteostasis and cause cytotoxicity in yeast (Choe et al., 2016; Yonashiro et al., 2016; Defenouillere et al., 2016). Whether this relates to the neurodegeneration caused by mutations in certain RQC genes in mammals (Chu et al., 2009; Ishimura et al., 2014) remains to be seen, as CAT-tailing is not known to occur in metazoans.

Mitochondrial dysfunction is another pathological hallmark of neurodegenerative disease (Johri and Beal, 2012; Coskun et al., 2012). In addition to producing ATP, mitochondria host a vast array of metabolic pathways and control processes such as Ca^2+^ homeostasis and apoptosis (Nunnari and Suomalainen, 2012). Mitochondrial dysfunction is profoundly implicated in major human diseases, from neurodegenerative and neuropsychiatric diseases to stroke, diabetes, and cancer (Sorrentino et al., 2018). Disease-associated proteins frequently interact with or enter into mitochondria and interfere with essential functions (Coskun et al., 2012). Moreover, environmental toxins can cause disease by targeting mitochondria (Sherer et al., 2002). How mitochondrial dysfunction arises and contributes to the often tissue- and cell type-specific pathologies remains enigmatic. While bioenergetic failure may partially explain the susceptibility of energy-demanding neuromuscular tissues, energy supplementation has been largely ineffective in treating diseases associated with mitochondrial dysfunction (Weber and Ernst, 2006), suggesting the involvement of other pathogenic events.

The co-presence of proteostasis failure and mitochondrial dysfunction as pathological hallmarks suggests that they are mechanistically linked. In the *Drosophila PINK1* model of PD with primary mitochondrial dysfunction, reduction of global protein synthesis to relieve burden on protein QC systems was beneficial (Liu and Lu, 2010). In a mouse model of Leigh syndrome caused by mitochondrial complex-I defect, inhibition of mTORC1, a regulator of mRNA translation, attenuated disease progression (Johnson et al., 2013). These studies strongly support an intimate connection between mitochondria and cytosolic translation, although the underlying mechanism is poorly defined.

Here we provide a molecular link between mitochondrial function and cytosolic translation/proteostatsis. Mitochondrial damage induces translational stalling of mitochondrial outer membrane (MOM)-associated *C-I30* mRNA by impairing the termination and ribosome-recycling factors eRF1 and ABCE1. Stalled ribosomes continue to add certain AAs to the C-terminus of C-I30, in a process analogous to CAT-tailing but with distinct features. C-I30 with C-terminal extension (CTE) inhibits oxidative phosphorylation (OxPhos) when assembled into C-I, and disrupts proteostasis when aggregating in the cytosol. Genetic manipulations of eRF1, ABCE1, or conserved CAT-tailing machinery prevent C-I30 CTE formation and rescue neuromuscular degeneration in *PINK1* flies. PINK1/Parkin mechanistically regulates the RQC and CTE processes. These results identify C-I30 and possibly ATP5a as the first endogenous metazoan substrates of the CTE process, revealing intimate connections between mitochondrial health, co-translational QC, and proteostasis in neuromuscular tissues.

## RESULTS

### A Novel Form of C-I30 in *PINK1* Neuromuscular Tissues

Human C-I is the largest enzyme of the respiratory chain, comprising 45 nuclear- or mitochondrial-encoded subunits, with C-I30 being part of the core assembly (Formosa et al., 2018). Mutations in *C-I30* cause OxPhos and neuromuscular defects in Leigh syndrome. *C-I30* mRNA is translationally repressed in the cytosol before being recruited to MOM and reactivated by PINK1 and Parkin (Gehrke et al., 2015), two factors linked to familial PD.

In muscle and brain tissues of *PINK1* flies, we detected canonical C-I30, which ran at ∼ 26 kD on SDS-PAGE gel, reflecting removal of the targeting sequence (MTS) from the preprotein. This was the only C-I30 band detected in WT flies. Intriguingly, an additional slower migrating band (C-I30-u) was detected in *PINK1* flies (Figure 1A). C-I30-u was detectable on high percentage (12-15%) SDS-PAGE gels running in the Tris/glycine/SDS but not MOPS buffer system (data not shown), explaining why it was not consistently observed in previous studies. Both C-I30 and C-I30-u were reduced in *PINK1* flies expressing independent *C-I30* RNAi transgenes (Figure 1B), demonstrating antibody specificity. C-I30-u was also observed in *parkin* flies (Figure 1C).

**Figure 1.**
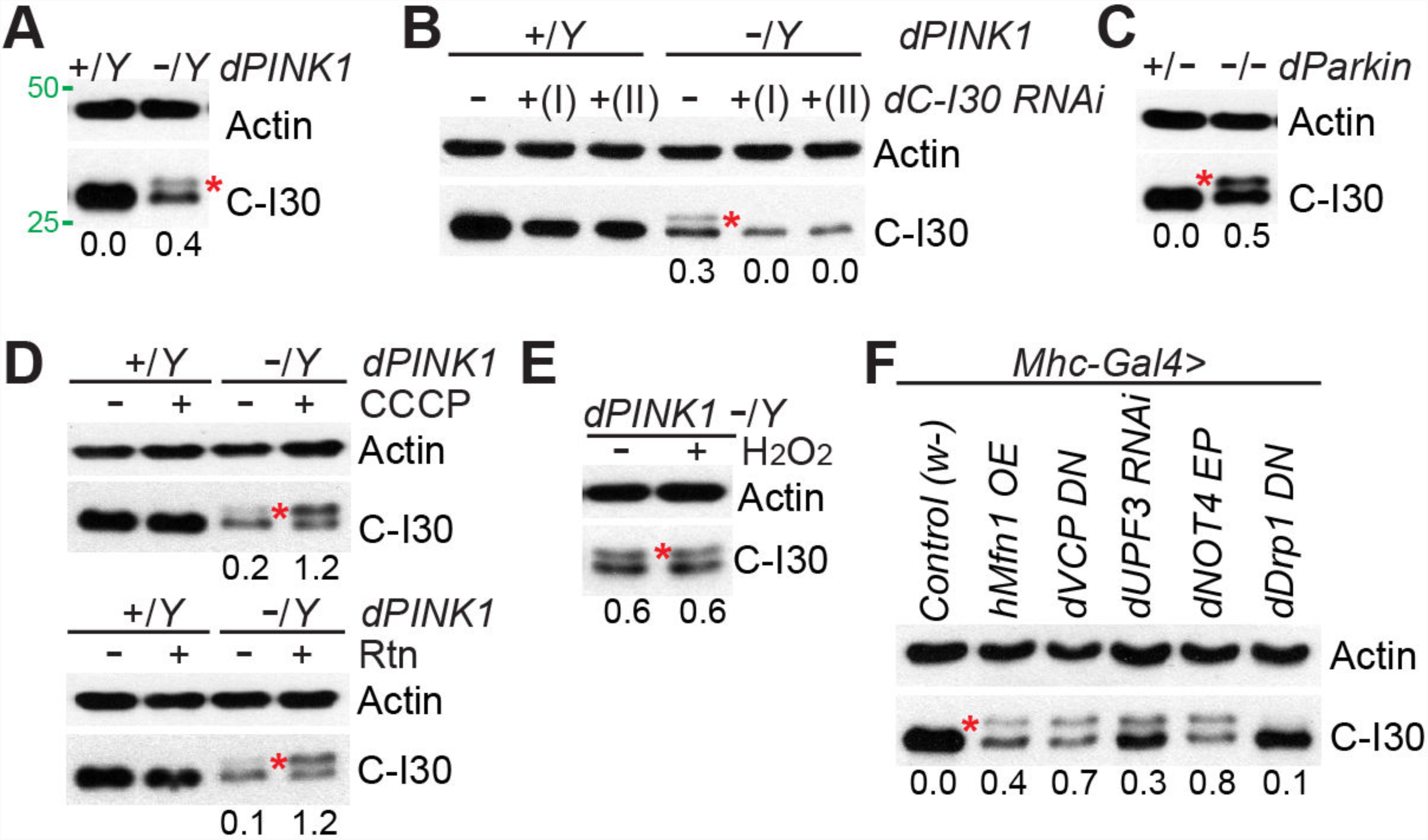
C-I30-u Formation in Mutant Flies Suffering Mitochondrial Insults. (A) Immunoblots of thoracic muscle extracts from *WT* (*dPINK1+/Y*) and *PINK1* (*dPINK1-/Y*) flies. * indicates C-I30-u. Values indicate ratio of C-I30-u/C-I30, and actin serves as loading control in this and subsequent Figures. (B) Immunoblots of muscle samples from *WT* and *PINK1* flies with C-I30 knocked down by 2 different *C-I30 RNAi* transgenes. (C) Immunoblots of muscle samples from *dParkin*^*1/+*^ and *dParkin*^*1/Δ 21*^ flies. (D) Immunoblots of muscle samples from *WT* and *PINK1* flies with or without CCCP or rotenone (Rtn) treatment. (E) Immunoblots of muscle samples from *PINK1* flies with or without H_2_O_2_ treatment. (F) Immunoblots of muscle samples from *Mhc-Gal4*-driven *UAS-hMfn1, dVCP RNAi, dUPF3 RNAi, dNOT4 EP* and *dDrp1 DN* transgenic flies. *w-*: WT control. See also Figure S1.

### C-I30-u Induction by Mitochondrial Stress

To probe how C-I30-u was formed, we pharmacologically treated *PINK1* flies. The C-I inhibitor rotenone, or the protonophore and mitochondrial uncoupler CCCP, caused a significant increase of C-I30-u relative to C-I30 (Figure 1D), but H_2_O_2_ had no obvious effect (Figure 1E). Knocking down subunit C of C-V also increased C-I30-u/C-I30 ratio, but surprisingly subunit α (bellweather, blw) knockdown showed opposite effect (Figure S1A). Previous studies showed that overexpression (OE) of the mitochondrial fusion GTPase hMfn1 (Ziviani et al., 2010), or a dominant-negative form of the AAA+ ATPase VCP (Kim et al., 2013), caused mitochondrial dysfunction and muscle degeneration (Figure S1B, C), phenocopying *PINK1* or *parkin*. These conditions also induced C-I30-u (Figure 1F). Although inhibiting the fission GTPase Drp1 has similar effect as hMfn1-OE on mitochondrial morphology (Ziviani et al., 2010), it did not obviously affect C-I30-u (Figure 1F), suggesting that the effect of hMfn1 on C-I30-u is independent of organelle fission/fusion. Thus specific types of insult induce C-I30-u.

### Translational Control of C-I30-u Formation

We explored the molecular basis of C-I30-u formation. One possibility is alternatively splicing. Our RT-PCR analysis failed to identify such mRNAs in *PINK1* mutant or in response to stress. Arguing against this possibility, C-I30-u still formed when a fully-spliced *C-I30* cDNA was expressed as a tagged transgene (C-I30-3xHA) in *PINK1* but not WT flies (Figure S2A). We note that even with the same transgene, C-I30-3xHA level was consistently lower in WT flies than *PINK1* flies, which have lower endogenous C-I30, suggesting some homeostatic regulation of C-I30 (Figure S2A). Moreover, when cDNA encoding Flag-tagged C-I30 was expressed in HeLa cells, C-I30-u was induced by either CCCP or the respiratory chain inhibitors Oligomycin and Antimycin A (Figure 2A; S2B), although a low level of tagged C-I30-u was detectable in HeLa cells without treatment (see explanation later). Phosphatase or ubiquitin protease failed to remove C-I30-u, suggesting that it is not formed by phosphorylation or ubiquitination (Figure 2B, C). C-I30-u was significantly higher in HeLa cells, which express negligible level of Parkin, compared to HeLa cells stably transfected with GFP-Parkin (Figure 2D), suggesting that Parkin restricts the formation of C-I30-u, the mechanism of which will be examined later.

**Figure 2.**
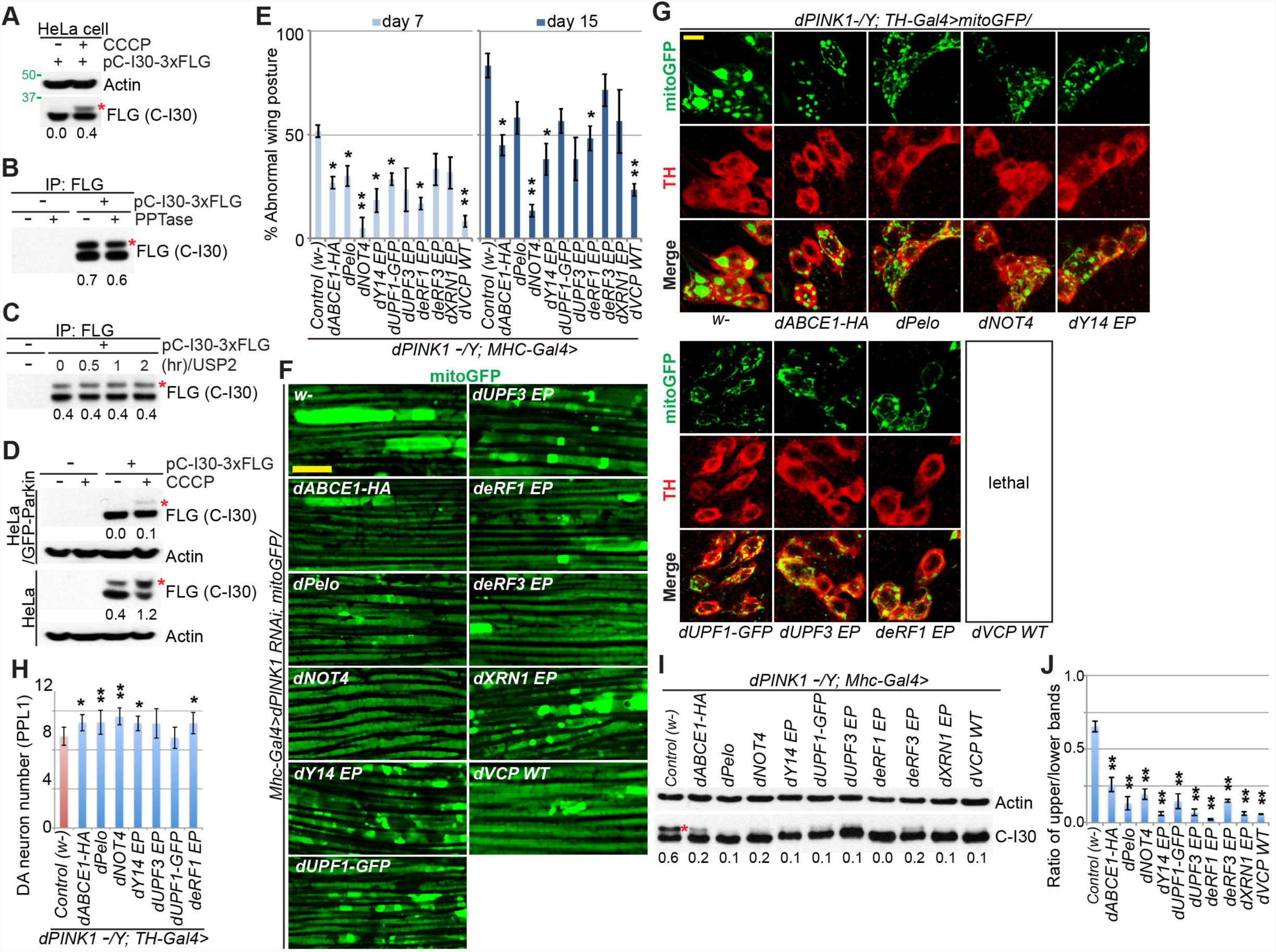
Co-Translational QC Genes Regulate C-I30-u Formation. (A) Immunoblots of lysates from HeLa cells transfected with 3xFLAG-tagged C-I30 cDNA and treated with 20 µM CCCP. (B, C) Immunoblots of PPTase-treated (B) or USP2-treated (C) C-I30-FLAG and C-I30-FLAG-u immunoprecipitated from HeLa cells. (D) Immunoblots of cell lysates from HeLa or GFP-Parkin/HeLa cells transfected with pCI-30-3xFLAG and treated with 20 µM CCCP. (E) Rescue of *PINK1* wing posture defect by co-translational QC factors. * *p*<0.05 and ** *p*<0.01 in SNK-test plus Bonferroni correction vs. *PINK1* group at day 7 or 15, respectively. (F) Removal of aberrant mitochondria by OE of co-translational QC factors in *dPINK1* RNAi fly muscle. (G, H) Rescue of *PINK1* DA neuron mitochondrial morphology (G) and number (H) by co-translational QC factors. H shows quantification of data shown in G. * *p*<0.05 and ** *p*<0.01 in SNK-test plus Bonferroni correction vs. *control (w-)*. (I) Immunoblots of muscle samples from *PINK1* flies expressing *UAS-dABCE1-HA, UAS-dPelo, UAS-NOT4-myc, dY14 EP, UAS-UPF1-GFP, dUPF3 EP, deRF1 EP, deRF3 EP, dXRN1 EP* and *UAS-dVCP* transgenes driven by *Mhc-Gal4*. *w-*: WT control. EP: enhancer P-element for OE. (J) Data quantification of I. ** *p*<0.01 in SNK-test plus Bonferroni correction vs. *control (w-)*. Error bars: s.d. from 3 independent assays and normalized with control, in this and all subsequent figures. Scale bar: 5µm in F, G, H. Mitochondrial morphology is monitored with a mito-GFP reporter. See also Figure S2.

Another possibility is that C-I30-u represents C-I30 preprotein. This is unlikely since unlike other proteins synthesized as preprotein in the cytosol and imported into mitochondria where MTS is removed, C-I30 engages in co-translational import on MOM (Gehrke et al., 2015). Thus, shortly after the MTS exiting the ribosome, it would be translocated by TOM/TIM and cleaved by the processing peptidases. Consistently, mass spectrometry (Mass Spec) analysis of C-I30-u failed to identify the MTS (see later). To further test this possibility, we mutated the MTS cleavage site in C-I30-Flag. This construct also produced two bands, with the lower band having a higher MW than WT C-I30-Flag, indicating blockage of MTS processing, and the upper band promoted by mitochondrial stress (Figure S2C). This ruled out C-I30-u being a preprotein.

### Co-translational QC Pathways Regulate C-I30-u Formation

We sought to gain insight into C-I30-u formation through systematic genetic analysis. Given that PINK1/Parkin regulates *C-I30* translation on MOM (Gehrke et al., 2015), we focused on genes in co-translational QC pathways, including the nonsense mediated decay (NMD), no-go decay, non-stop decay, and RQC pathways (Brandman and Hegde, 2016). Although these pathways target aberrant mRNAs with distinct features, they are mechanistically related and share certain components (Brandman and Hegde, 2016). We used the *PINK1* wing posture phenotype, caused by mitochondrial and muscle degeneration, as readout of genetic interaction. Remarkably, OE of co-translational QC factors, with the OE effect verified for certain key factors (Figure S2D), effectively rescued *PINK1* phenotypes in the muscle (Figure 2E, F) and disease-relevant DA neurons (Figure 2G, H). Correlating with the phenotypic rescue, C-I30-u level was greatly reduced (Figure 2I, J). Loss-of-function of the co-translational QC genes in *PINK1* mutant showed little effect on C-I30-u as compared to *PINK1* mutant alone (Figure S2E, F), likely because they work in the same pathway to regulate C-I30-u. However, there are exceptions, as shown later.

Consistent with the co-translational QC genes working in the same pathway as PINK1 to control C-I30-u expression and mitochondrial function, muscle-specific knockdown of *UPF3*, a key NMD factor (Leeds et al., 1992), was sufficient to phenocopy *PINK1*. Similarly, muscle-specific expression of dominant-negative *VCP*, a key RQC factor (Brandman et al., 2012), or overexpression of NOT4, a protein implicated in the co-translational QC of C-I30 (Wu et al., 2018), also phenocopied *PINK1* (Figure 1F; S1B, C).

One exception to the general trend of co-translational QC gene interaction with PINK1 is Clbn (Bi et al., 2005), the fly homologue of yeast RQC2/Tae2. RQC2/Tae2 recruits charged Ala- and Thr-tRNAs to stalled ribosomes and is required for CAT-tailing in yeast (Shen et al., 2015). In CLIP assays, we found that Clbn binds to fly Ala- and Thr-tRNAs, and to a lesser extent, Ser-tRNA (Figure 3A). Complete loss of *clbn* had no obvious effect under normal conditions, as we reported before (Wang et al., 2013b). When introduced into *PINK1* mutant, complete loss of *clbn* abolished C-I30-u formation, whereas 50% loss had no obvious effect (Figure 3B). Complete loss of *clbn* rescued the wing posture (Figure 3C), ATP production (Figure S3A), mitochondrial morphology (Figure 3D; S3B), and DA neuron loss (Figure 3E) phenotypes of *PINK1*.

**Figure 3.**
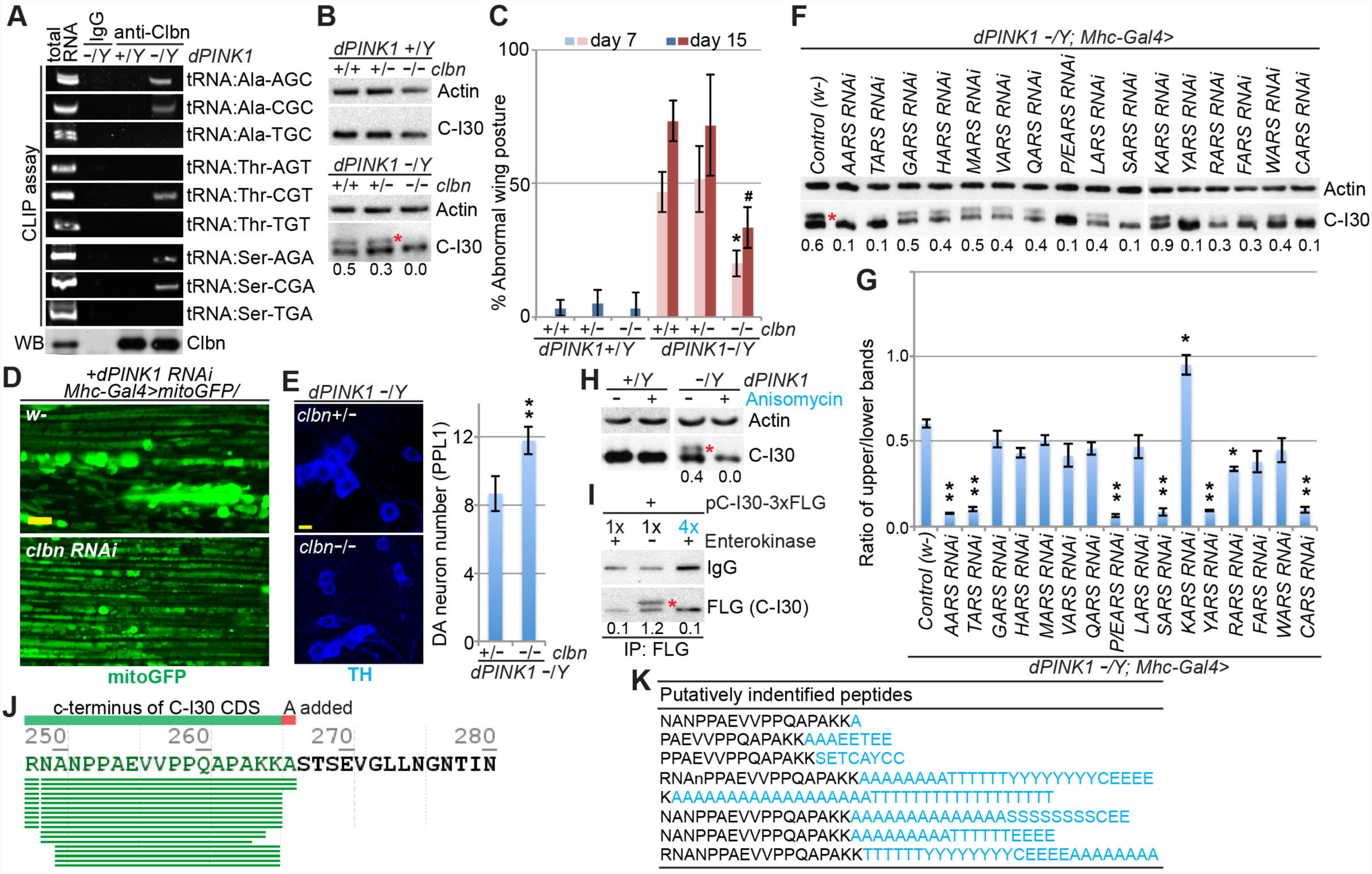
C-I30-u Behaves as a CTE Form of C-I30. (A) Cross-link immunoprecipitation (CLIP) assay showing binding of specific tRNAs by Clbn in *PINK1* fly muscle tissue. tRNAs cross-linked to Clbn were analyzed by RT-PCR. (B) Immunoblots of muscle samples from *WT* and *PINK1* flies with loss of 1 or 2 copies of *clbn.* (C) Effect of *clbn* on wing posture in *WT* and *PINK1* condition. * or #, *p*<0.05 in SNK-test plus Bonferroni correction vs. *PINK1* at day 7 or 15, respectively. (D) Effect of *clbn* RNAi on mitochondrial morphology in *dPINK1* RNAi fly muscle. Mitochondrial morphology is monitored with a mito-GFP reporter. (E) Rescue of DA neuron number by *clbn* in *PINK1*. ** *p*<0.01 in two-tailed Student’s *t*-test. (F, G) Immunoblots of muscle samples from *PINK1* flies expressing various *AARS* RNAi transgenes driven by *Mhc-Gal4* (F). Data quantification is shown in G. * *p*<0.05 and ** *p*<0.01 in SNK-test plus Bonferroni correction vs. *control (w-)*. (H) Immunoblots of muscle samples from *WT* and *PINK1* flies treated with anisomycin. (I) Immunoblots of enterokinase (EK)-treated C-I30-FLAG and C-I30-FLAG-u purified from HeLa cells. After digestion, the remaining protein showed reduced Flag signal, presumably due to the removal of one residue from the epitope. 1x and 4x of the resulting products were loaded to match signal in the uncut sample. (J) C-terminal peptides identified from generic database searches of fly C-I30-u Mass Spec data (see Table S1). SEVGLLNGNA indicates read-through sequence. (K) Putatively identified longer CTE peptides from customized pool-based searches. aa in black: C-I30 CDS; aa in blue: CTE (see Table S2). Scale bar: 5µm in D, E. See also Figure S3.

The above results supported C-I30-u formation by a CAT-tailing-like process. To test this further, we examined the requirement for Ala- or Thr-tRNA synthetase (AARS or TARS). Partial inhibition of AARS or TARS preferentially reduced C-I30-u formation (Figure 3F, G) and mitochondrial aggregation caused by *PINK1* inhibition (Figure S3C). SARS, YARS, CARS, or E/PARS had similar effect (Figure 3F, G; S3C). The specificity of the ARS effect was demonstrated by lack of effect of the rest of the ARSs tested, with the exception of KARS, whose RNAi moderately increased C-I30-u (Figure 3F, G). Anisomycin, which targets the peptidyl-transferase center and can inhibit CAT-tailing *in vitro* (Osuna et al., 2017), inhibited C-I30-u formation in fly muscle (Figure 3H; S3D). These data indicate that C-I30-u is formed through the addition of A, T, and possibly other select AAs in a CAT-tailing like process.

### C-I30-u is a CTE Form of C-I30

To further characterize C-I30-u, we made use of C-I30 constructs containing epitope tags allowing C-terminal analysis. Transfected HeLa cells were treated with CCCP to induce C-I30-u formation. In the C-I30-Flag-AK construct, a 3xFlag tag was inserted 2 codons (encoding AK) before the *UAG* stop codon. With this construct, we observed CCCP induction of C-I30-Flag-u (Figure 2A). Cleavage of Flag with enterokinase removed C-I30-Flag-u (Figure 3I), consistent with C-I30-Flag-u having CTE. Although we designed our constructs with the initial thinking that the K at the very end might play a role in CTE, deleting K or adding another K had no obvious effect on C-I30-u formation (Figure S3E). We made two other constructs, C-I30-TEV-Flag and C-I30-Flag-TEV, with a tobacco etch virus (TEV) protease-cleavage site inserted before or after Flag. Both constructs exhibited C-I30-u induction by CCCP (Figure S3F). With C-I30-TEV-Flag, we could trace the TEV protease-released CTE with anti-Flag. Two Flag-positive fragments (Flag alone and Flag with tail) were detected, again supporting C-I30-TEV-Flag-u having CTEs (Figure S3G).

To determine the nature of the CTE in C-I30-u, we performed Mass Spec of C-I30-u purified from *PINK1* flies. We took advantage of the fact that C-I30-u is less soluble than C-I30. By enriching for insoluble proteins followed by denaturing IP, we obtained relatively pure C-I30-u (Figure S3H), which was subjected to gel purification, in-gel partial digestion with trypsin, and Mass Spec of released peptides. No strong signals for peptides containing MTS, or PTMs such as phosphorylation or ubiquitination were found. We also did not find evidence of read-through after the stop codon. Based on *in vivo* ARS-dependency of C-I30-u, we hypothesized that A/T/S/Y/C/E/P might be the preferred AAs in the CTE, and built custom library with that in mind. Mass Spec data supported that notion. For example, search of tandem mass spectra against the databases frequently identified the peptide with A added to the very C-terminal fragment of C-I30 (RNANPPAEVVPPQAPAKK.A), suggesting that A is more preferred at the first position of the CTE (Figure 3J, K; Table S1). Since C-I30-u ran as a rather discrete band on SDS-PAGE with an estimated MW ∼ 3-4 kD larger than C-I30, we searched for peptides with longer CTE against customized databases. Due to size limitation of the database that could be used in our Mass Spec analysis, we were not able to exhaust searching all possible AA combinations of the 3-4 kD CTE. Nevertheless, our searches led to the putative identification of longer CTEs containing A, T, and the other preferred AAs (Figure 3K; Table S2). Peptide spectra produced by collision-induced dissociation were identified and manually examined (Figure S3I). The shorter peptides we identified might result from cleavage of the CTEs by trypsin, or length-heterogeneity of the CTEs hidden by the discrete banding pattern of C-I30-u on SDS-PAGE. As controls, we performed Mass Spec of C-I30 purified from WT flies, but did not find CTEs (data not shown). We also generated databases of peptides composed of AAs whose ARSs knockdown did not affect C-I30-u. Such searches came up mostly negative. Thus, C-I30 CTE in *Drosophila* is analogous to CAT-tailing in yeast, but with distinct features.

### Stress-Induced Defect in Translation Termination Triggers C-I30-u Formation

A unique feature of C-I30 CTE is that AAs are added to full-length C-I30, instead of incomplete NPCs. We wondered if this is because mitochondrial stress affects translation termination at the normal stop codon. We found that CCCP reduced levels of eRF1 and ABCE1, but not eRF3, in HeLa cells (Figure 4A). Since eRF1 and ABCE1 form a complex (Preis et al., 2014), mitochondrial damage may target one or both components of this complex. Correlating with inhibition of C-I30-u formation by Parkin (Figure 2D), Parkin blocked the CCCP effect on eRF1 and ABCE1 (Figure 4B). Parkin also promoted ABCE1/eRF1 and ABCE1/Pelo interaction (Figure S4A), suggesting that it normally ensures the fidelity of translation termination and/or ribosome recycling.

**Figure 4.**
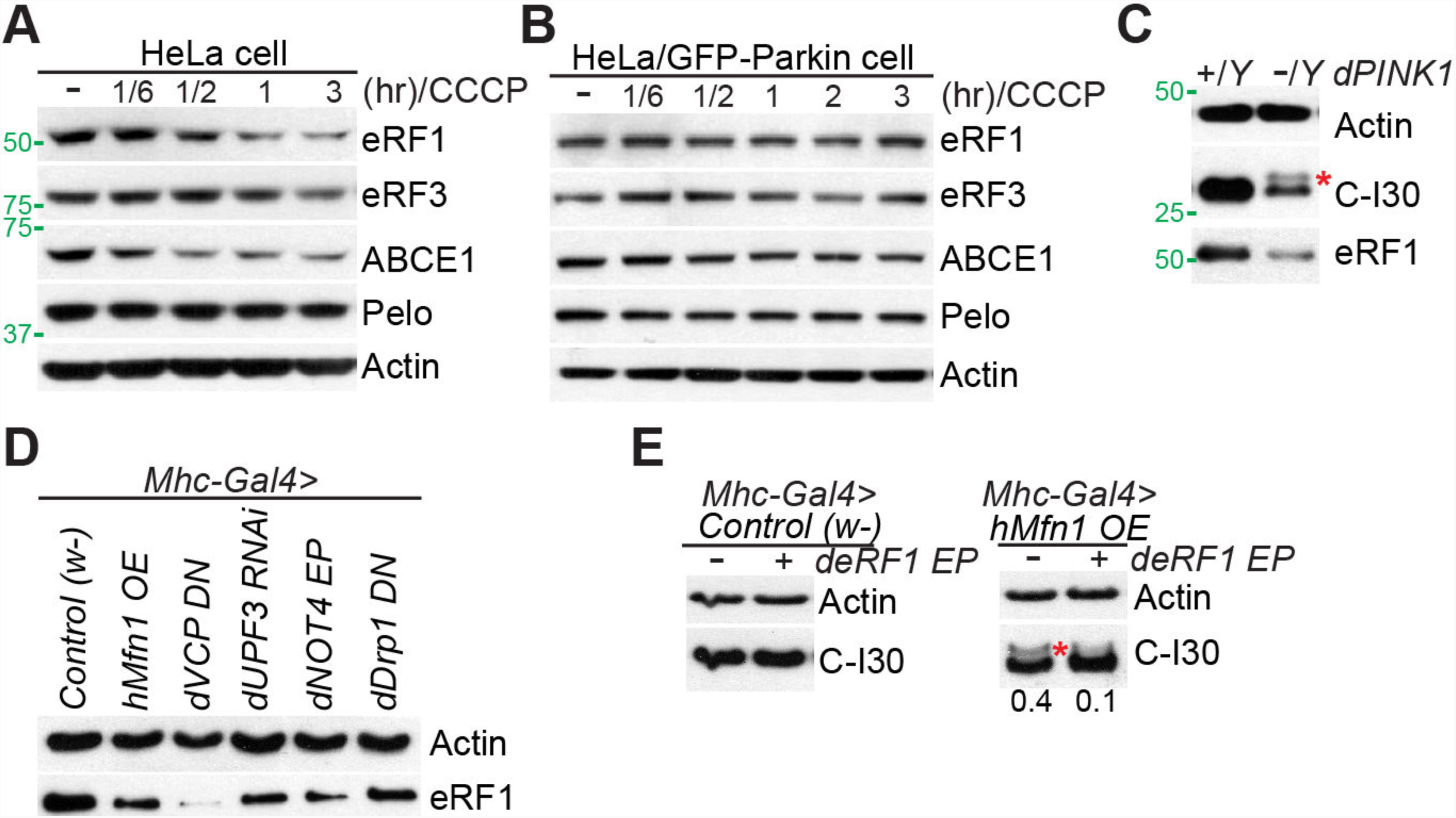
eRF1 Regulates C-I30-u Formation. (A, B) Immunoblots of indicated proteins in CCCP-treated HeLa cells (A), or HeLa/GFP-Parkin cells (B). (C) Immunoblots of eRF1 in muscle sample of *WT* and *PINK1* flies. Same blot as in Figure 1A is used here. (D) Immunoblots of eRF1 in muscle samples from WT flies expressing *hMfn1-OE, dVCP RNAi, dUPF3 RNAi, dNOT4-OE*, and *dDrp1-DN* transgenes driven by *Mhc-Gal4*. *w-*: WT control. (E) Immunoblots showing removal of C-I30-u by eRF1-OE in *Mhc-Gal4>hMfn1* flies. See also Figure S4.

We next tested the *in vivo* relevance of the cell culture studies. The level of full-length eRF1 was reduced in *PINK1* muscle tissue (Figure 4C). Due to the lack of antibodies, we could not examine fly ABCE1 and eRF3. eRF1 level was also reduced in UPF3-RNAi, VCP-DN, NOT4, or hMfn1 OE flies (Figure 4D), which phenocopied *PINK1* in C-I30-u formation and muscle degeneration. Intriguingly, concomitant with the reduction of full-length eRF1, a smaller-sized eRF1 appeared (Figure S4B), suggesting that eRF1 might be processed in these mutant conditions.

We next used eRF1-OE to assess the causal role of eRF1 in C-I30-u formation. eRF1-OE reduced C-I30-u in *PINK1* (Figure 2I) or *hMfn*-OE flies (Figure 4E), and rescued their mitochondrial and muscle phenotypes (Figure 2E, 2F, S4C). Moreover, eRF1-OE rescued the mitochondrial and DA neuron degeneration in *PINK1* flies (Figure 2G, H). ABCE1-OE had similar effect (Figure 2E-H, I). Supporting C-I30 being a critical target mediating eRF1 and ABCE1 effects, C-I30 RNAi blocked the rescue of *PINK1* by eRF1-, ABCE1-, or UPF3-OE, although C-I30 RNAi alone did not cause toxicity or modify *PINK1* phenotypes (Figure S4D, E).

These results establish impaired translation termination due to eRF1/ABCE1 alteration as a cause of mitochondrial stress-induced translational stalling and ensuing RQC. We term this phenomenon <u>mi</u>tochondrial <u>s</u>tress-induced translational <u>term</u>ination impairment a<u>n</u>d protein c<u>a</u>rboxyl terminal <u>e</u>xtension (MISTERMINATE).

### C-I30-u Assembly into Respiratory Chain Inhibits Energetics and Cellular Survival

The strict correlation between C-I30-u abundance and disease severity *in vivo* suggests that C-I30-u may interfere with mitochondrial function and neuromuscular integrity. In the recently solved C-I structure, the C-terminus of C-I30 was found in the vicinity of NDUFS1, NDUFS4, NDUFA5, and NDUFA7, but the last 14-aa (SLKLEAGDKKPDAK) was not included due to poor electron density (Gu et al., 2016). The very C-terminus of C-I30 may exist in a dynamic state and be regulatory. If so, assembly of C-I30-u into C-I may alter C-I function. We found that in contrast to mito-dsRed, which was synthesized in the cytosol (cyto) and imported into mitochondria (mito) and thus present in both the cyto and mito fractions, C-I30 was found exclusively in the mito fraction (Figure S4F). Moreover, using a C-I30 construct containing N-term HA and C-term Flag, we found mito-associated C-I30 intermediate species positive for HA but not Flag, likely representing partially synthesized C-I30 (Figure S4G). Together, these data further supported the co-translational import of C-I30. Using 2-D gel, with blue native gel electrophoresis (BNG) being the 1st dimension to separate large mitochondrial complexes, and SDS-PAGE the 2nd dimension to separate constituents of the complexes, we found both C-I30 and C-I30-u in C-I of *PINK1* muscle tissues (Figure 5A) or human cells (Figure S5A). The complexes containing C-I30 and C-I30-u were verified to be C-I and C-I-associated super-complex based on co-migration of C-I marker (Figure 5A).

**Figure 5.**
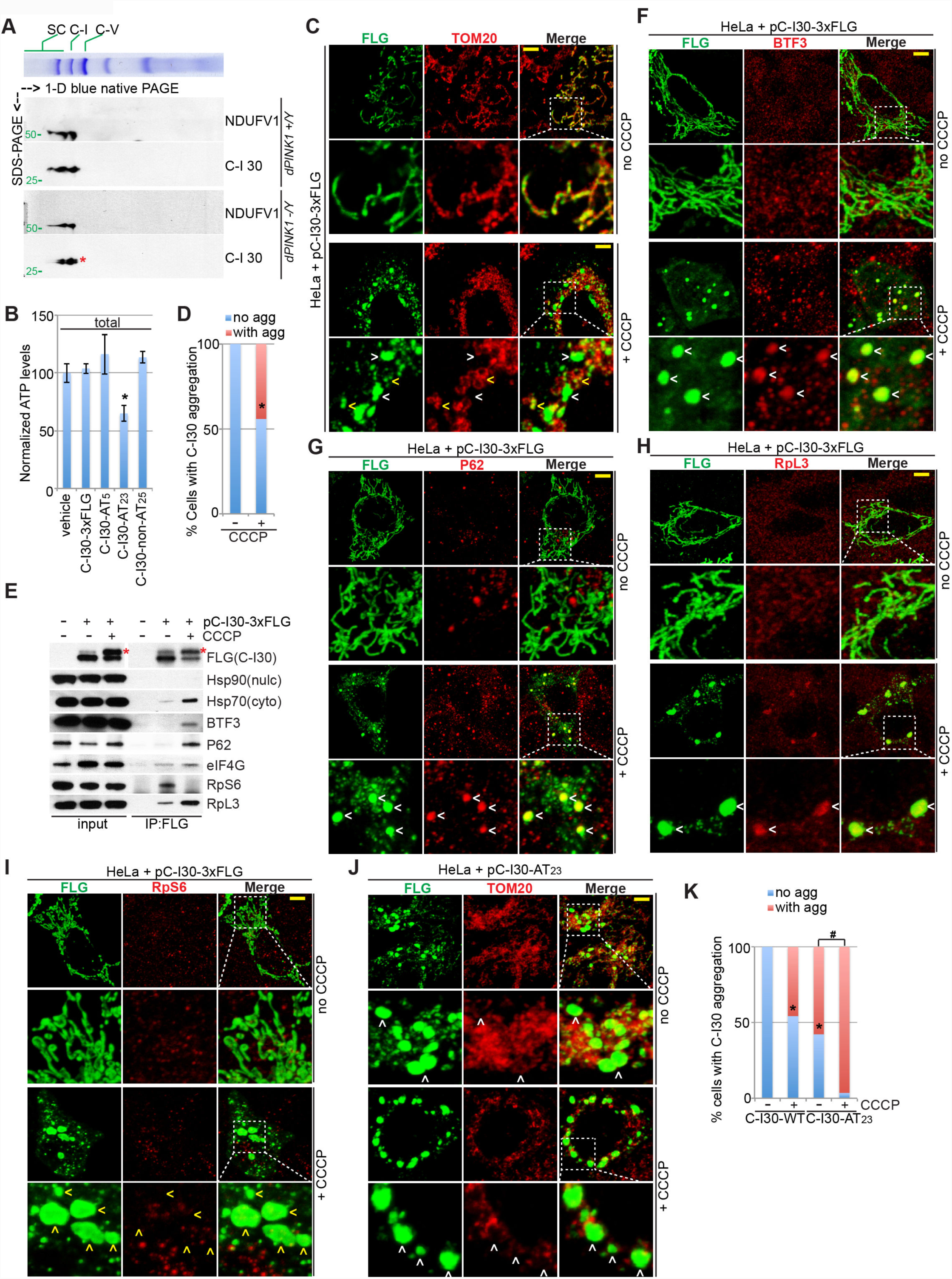
C-I30-u can be Assembled into C-I and Form Cytosolic Aggregates. (A) Immunoblots of C-I30 by 2D gel showing C-I30-u assembly into the RCC complex in purified mitochondria from *PINK1* flies. C-I30 and C-I30-u are present in complex-I (C-I) and supercomplexes (SC), the identities of which are verified by the co-migration of another C-I marker (NDUFV1). (B) ATP measurement in HeLa cells expressing C-I30-CAT-Tail constructs. * *p*<0.05; in SNK-test plus Bonferroni correction vs. *control (w-)*. (C) Immunostaining showing C-I30 aggregates in CCCP-treated HeLa cells. Scale bar, 3µm. Arrowheads: aggregates outside (white) or inside (yellow) mitochondria. (D) Qualification of data shown in D, with % of cells with C-I30 aggregates indicated. *, *p*<0.05; *Chi-*squared test. (E) Immunoblots of FLAG IP from HeLa cells transfected with C-I30-FLAG, with or without CCCP treatment. (F-I) Immunostaining of HeLa cells transfected with C-I30-FLAG and treated with CCCP. Scale bar, 3µm. Arrowheads, white: aggregates colocalizing with BTF3 (F), P62 (G), or RpL3 (H); yellow: no RpS6 colocalization (I). (J) Immunostaining of HeLa cells transfected with C-I30-Flag-AT_23_, and with or without CCCP. Arrowheads: aggregates outside mitochondria. (K) Qualification of data shown in K, with % of cells with aggregates indicated. *, *p*<0.05; *Chi-* squared test. #, *p*<0.05; *Chi-*squared test, compared to C-I30-Flag-AT_23_ non-treated group. More than 100 transfected cells in each group were quantified in D and K. Scale bar, 3µm in C, F-I, J. See also Figure S5.

Since the removal of C-I30-u in *PINK1;clbn* flies (Figure 3B) correlated with increased ATP production compared to *PINK1* flies (Figure S3A), assembly of C-I30-u into C-I might inhibit C-I activity. To further test this possibility, we made a C-I30-u mimetic (C-I30-CAT-tail) containing C-term (AT)_23_ repeats. Expression of C-I30-CAT-tail reduced ATP production and C-I activity compared to C-I30-WT, C-I30 with shorter AT tail, or C-I30 with similar length but non-AT tail (Figure S5B-D; 5B). Moreover, mitochondria from C-I30-Flag transfected cells also showed mildly reduced C-I activity and ATP production compared to control cells (Figure S5C, D). Thus C-I30 has very stringent requirement for its C-terminus such that even the Flag tag partially interferes with its function and causes mitochondrial stress. This may explain the low level C-I30-Flag-u formation in non-treated HeLa cells.

### C-I30-u Forms Aggregates and Alters Proteostasis in a CTE-Dependent Manner

When allowed to persist, CAT-tailed GFP formed aggregate in the cytosol of yeast cells (Choe et al., 2016). To test whether C-I30-u forms aggregates in mammalian cells, we examined the localization of C-I30-Flag. C-I30-Flag transfected cells exhibited a Flag localization pattern similar to that of the MOM protein TOM20 under normal condition (Figure 5C), consistent with IMM localization of C-I30. Concomitant with the induction of C-I30-Flag-u by CCCP, prominent Flag+ aggregates formed (Figure 5C, D). CCCP did not induce aggregation of GFP-Flag (Figure S5E). Aggregates formed by endogenous C-I30 could also be observed in HeLa cell treated with CCCP (Figure S5F). While smaller aggregates tended to co-localize with mitochondria, larger aggregates were mainly in the cytosol (Figure 5C). The larger aggregates are likely cytosolic assemblies of C-I30-u, which was released from MOM where it was synthesized. In HeLa/GFP-Parkin cells where C-I30-u was restricted to a lower level, C-I30 aggregation was much less pronounced (Figure S5G, H).

We next examined proteins associated with C-I30-u aggregates to assess the state of proteostasis. Using cross-linking followed by fractionation to separate soluble proteins from insoluble aggregates, we enriched for CCCP-induced C-I30-u and then performed co-IP. Chaperones, e.g. nascent peptide-associated complex (NAC) subunit BTF3, Hsp70, and QC factors (p62) showed increased association with C-I30-u (Figure 5E). NAC normally binds to ribosomes to promote NPC folding, but under stress it may move to protein aggregates and function as chaperone (Preissler and Deuerling, 2012). We also found metastable proteins, e.g. eIF4G, in the aggregates (Olzscha et al., 2011). Consistent with C-I30-u being synthesized on split 60S subunit, 60S ribosomal proteins showed preferential association with C-I30-u (Figure 5E). Confocal microscopy validated localization of these factors in C-I30-u aggregates (Figure 5F-I). Further supporting C-I30-u forming aggregates through the CTE, C-I30-(AT)_23_ formed aggregates even in the absence of mitochondrial stress (Figure 5J, K, S5B), whereas WT C-I30-Flag formed aggregates to a much lesser extent and only after CCCP treatment (Figure S5I). We found that C-I30-u induction by CCCP increased the level of p-eIF2α, a marker of integrated stress response (Pakos-Zebrucka et al., 2016), and activated the Hippo/LATS/YAP signaling pathway known to be regulated by proteotoxicity caused by defective ribosomal products (Meriin et al., 2018), indicating that CTE-mediated aggregation perturbs proteostasis (Figure S5J).

In muscle tissue of *PINK1* but not wild type flies, endogenous C-I30 formed aggregates (Figure 6A). In DA neurons, transgenic C-I30-HA also formed aggregates specifically in *PINK1* flies (Figure 6B). Removal of *clbn* dramatically suppressed C-I30 aggregation (Figure 6C). Together with the data showing removal of C-I30-u in *PINK1;clbn* double mutant, these results support that aggregation is attributable to C-I30-u. Consistently, OE of eRF1 or ABCE1, knockdown of ARSs involved in CTE, or treating cells with anisomycin, conditions that inhibited C-I30-u formation, all prevented C-I30 aggregation (Figure 6D, E). Knockdown of eRF1, ABCE1, and the ARSs not involved in CTE, or *clbn*-OE did not further modify C-I30 aggregation (Figure S6A). Anisomycin also removed C-I30 aggregates in *VCP DN, hMfn1-OE*, and *UPF3 RNAi* fly muscle (Figure S6B, C). Thus C-I30 with CTE is prone to aggregation and causes proteostasis failure if not adequately removed.

**Figure 6.**
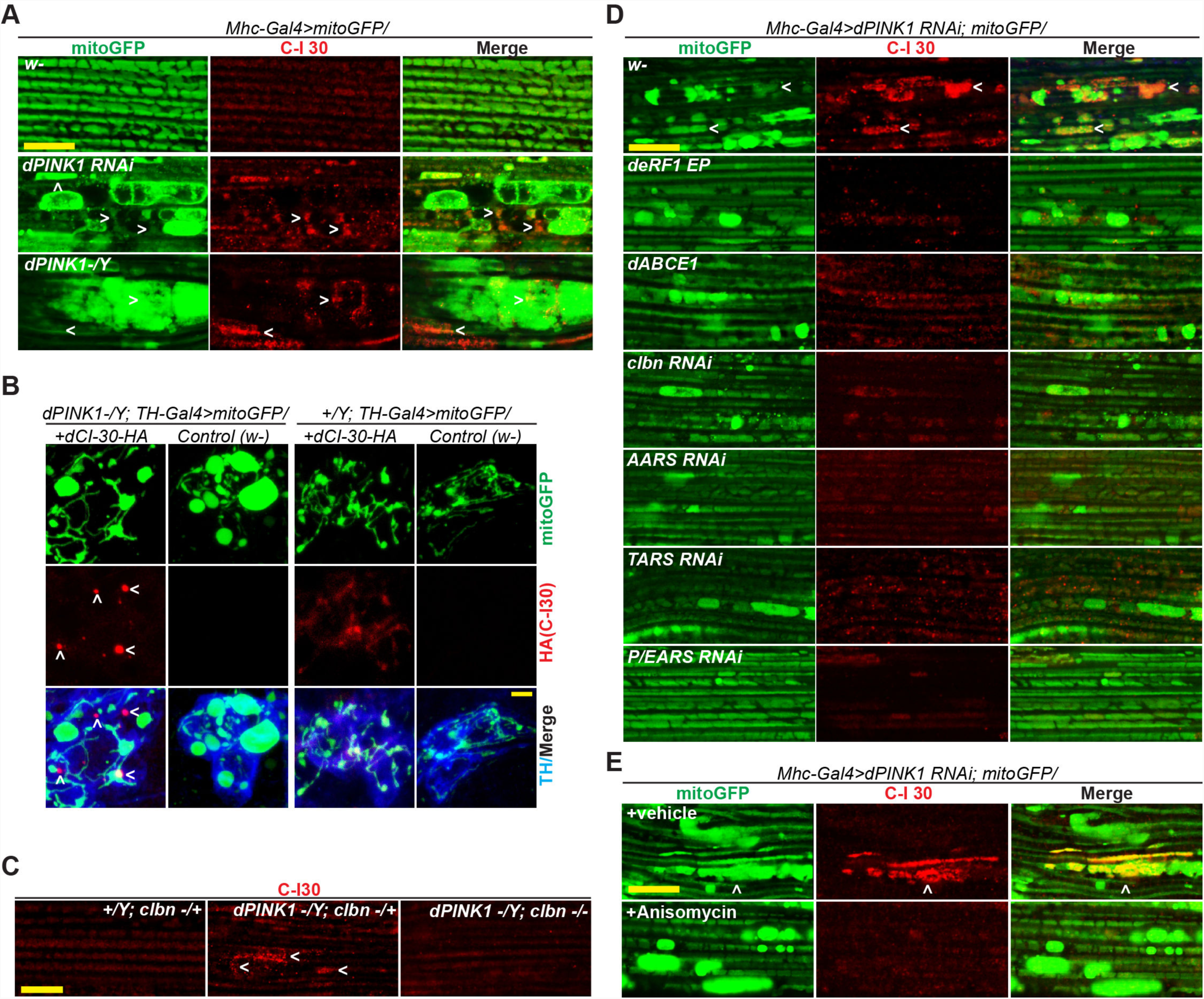
C-I30-u Forms Aggregates in Fly Neuromuscular Tissues. (A) Immunostaining showing C-I30 aggregates in *PINK1* muscle. (B) Immunostaining showing C-I30-HA aggregates in *PINK1* DA neurons expressing a C-I30-HA transgene. (C) Immunostaining showing effect of *clbn* on C-I30 aggregation in *PINK1* muscle. (D) Immunostaining showing effect of various genetic manipulations on mitochondrial morphology and C-I30 aggregation in *PINK1* muscle. (E) Immunostaining showing effect of anisomycin on C-I30 aggregation in *Mhc>PINK1 RNAi* fly muscle. Due to the acute nature of treatment, mitochondrial morphology is not rescued. Scale bars, 5µm. Arrowheads: CI-30 aggregates. Mitochondrial morphology is monitored with a mito-GFP reporter. See also Figure S6.

### Mechanisms of C-I30-u Formation in Mammalian Cells

The induction of C-I30-u and C-I30 aggregation by mitochondrial stress in mammalian cells offered us an opportunity to study the mechanisms involved. Both *C-I30* and *ATP5a* mRNAs are translated on MOM in mammalian cells (Gehrke et al., 2015). We found that endogenous C-I30 and ATP5a both underwent CTE (Figure 7A) and aggregation (Figure S7A) upon CCCP treatment of HeLa cells. As in *PINK1* flies, anisomycin could reduce the CTE of the endogenous proteins, and the CTE and aggregation of transfected C-I30-Flag (Figure 7B, C; S7D). CTE and aggregation of ATP5a were also seen in *PINK1* flies (Figure S7B, C).

**Figure 7.**
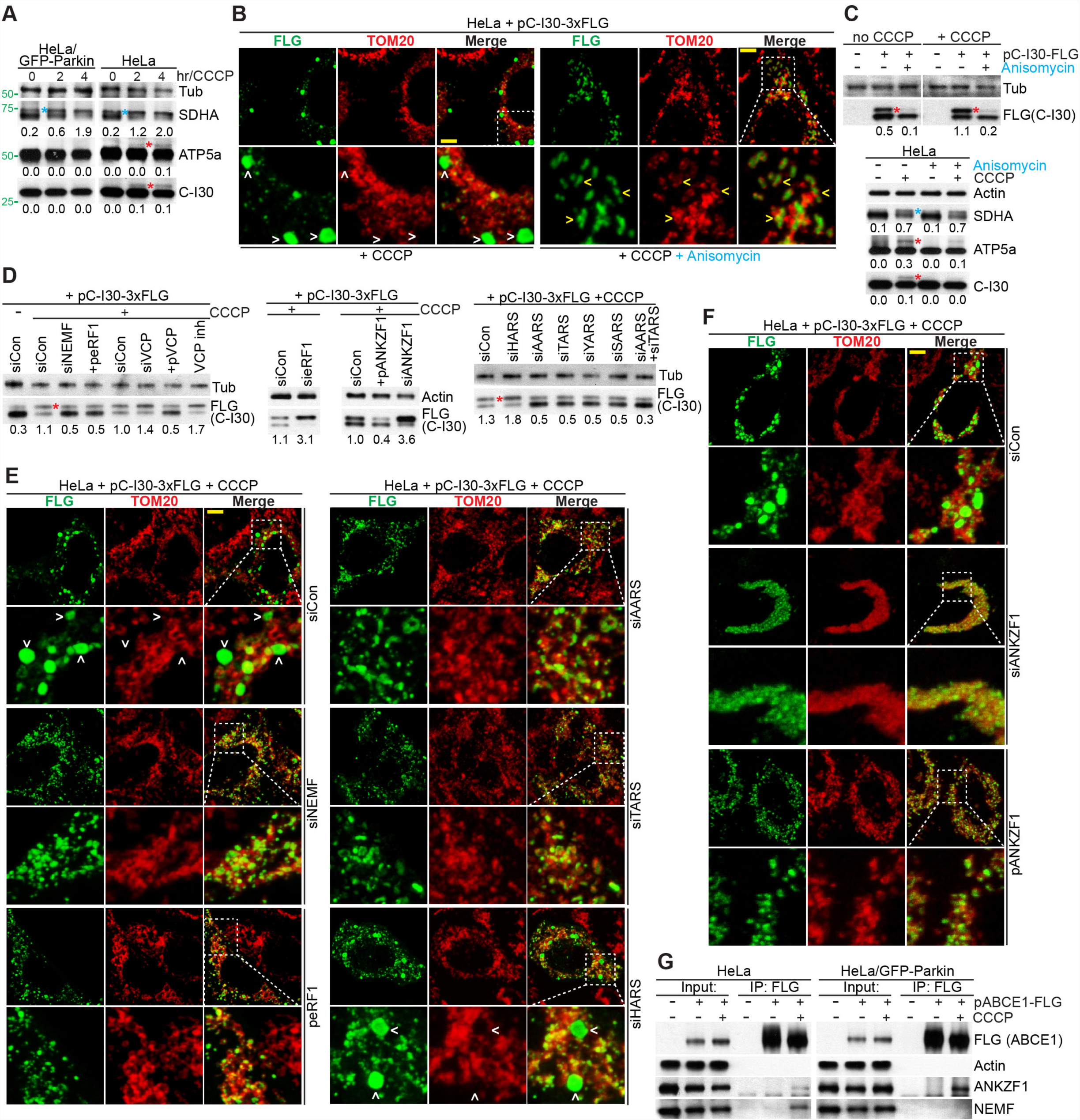
Mechanism of C-I30-u Formation and Aggregation in Human Cells. (A) Immunoblots showing CCCP-induced CTE forms (red *) of C-I30 and ATP5a in HeLa cells but not HeLa/GFP-Parkin cells. Blue * indicate preprotein of SDHA that accumulates in both HeLa and HeLa/GFP-Parkin cells upon CCCP treatment. (B) Immunostaining of HeLa cells transfected with C-I30-Flag and treated with CCCP or CCCP+anisomycin. Arrowheads: aggregates outside (white) or inside (yellow) mitochondria. (C) Immunoblots of showing effect of anisomycin treatment on the CTE forms of endogenous ATP5a and C-I30 or C-I30-Flag in HeLa cell. (D) Immunoblots showing effects of the various genetic manipulations or VCP inhibitor treatment on C-I30-Flag-u formation. (E) Immunostaining of HeLa cells co-transfected with C-I30-FLAG and various modifiers, showing effects on CCCP-induced FLAG+ aggregates. Arrowheads: aggregates. (F) Immunostaining of HeLa cells showing effects of ANKZF1 RNAi or OE on C-I30-Flag aggregation. (G) Immunoblots showing effect of PINK1 on ANKZF1 and NEMF interaction with ABCE1 in co-IP assays. Scale bar, 3µm in B, E, F. See also Figure S7.

We next tested the molecular players involved. RNAi of NEMF, the mammalian homologue of Tae2/Clbn, reduced C-I30-u formation and C-I30 aggregation (Figure 7D, E; S7F). eRF1- or ABCE1-OE also reduced C-I30-u level and C-I30 aggregation (Figure 7C, E; S7E, F). VCP negatively regulated C-I30-u, as its OE reduced, whereas inhibition by RNAi or a small molecule inhibitor increased C-I30-u level (Figure 7D). Consistent with select AAs being chosen for CTE, knockdown of A/T/Y/SARS, but not HARS, significantly reduced C-I30-u level and aggregation (Figure 7D, E, S7F). In the case of C-I30-AT_23_, inhibition of CTE also reduced its degree of aggregation (Figure S7G, H), suggesting that even without external stress, C-I30-AT_23_ already underwent additional CTE, possibly due to its intrinsic toxicity, and the additional CTE further contributed to aggregation.

Recently, Vms1/ANKZF1 was shown to protect mitochondria by antagonizing the CTE machinery (Izawa et al., 2017), at least in part by cleaving the peptidyl-tRNA bond (Verma et al., 2018). We tested the role of ANKZF1 in C-I30-u formation. ANKZF1 RNAi promoted, whereas its OE inhibited C-I30-u formation (Figure 7D). Strikingly, ANKZF1 RNAi resulted in C-I30 and C-I30-u largely accumulating on MOM, without cytosolic aggregation (Figure 7F), suggesting that ANKZF1 is normally involved in releasing C-I30-u from MOM into the cytosol. This result further supported MOM as site of C-I30/C-I30-u synthesis.

We also found that ANKZF1 and NEMF interacted with ABCE1 in HeLa cells, and that Parkin strengthened ABCE1/ANKZF1 but weakened ABCE1/NEMF interactions (Figure 7G). Parkin also promoted recruitment of ANKZF1 and eRF1, but inhibited that of NEMF, to MOM-associated 60S ribosomes in CCCP-treated cells (Figure S7I). Moreover, C-I30-u was specifically found on MOM-associated 60S ribosomes in HeLa cells (Figure S7I). These results further supported C-I30-u being made by a CTE process, and began to reveal upstream regulatory mechanisms. Finally, we tested the relevance of our findings to human disease. We found increased C-I30-u and ATP5a-u in aged *PINK1* and *Parkin* patient fibroblasts compared to fibroblasts from matched control subjects (Figure S7J).

## DISCUSSION

Timely removal of aberrant proteins is critical for cellular health. A number of major human diseases are associated with defective protein QC, resulting in disease-defining protein aggregates. The mechanism of protein aggregate formation has not been clearly defined. Our results demonstrate that defects in co-translational QC can lead to aberrant protein accumulation, aggregation, and proteotoxicity *in vivo* in settings relevant to neurodegenerative disease. We show that mitochondrial dysfunction can impair translation termination fidelity and impinge on co-translational QC, mechanistically linking mitochondrial dysfunction with proteostasis failure, two cardinal features of neurodegenerative disease.

### C-I30 and Possibly ATP5a are Endogenous Metazoan Proteins Modified by a CTE Process

Artificial templates have been instrumental in previous RQC studies. Before this study, CAT-tailing was observed in yeast on non-stop reporter proteins, where CAT-tails were heterogeneous in length, and composed exclusively of A and T (Shen et al., 2015). CAT-tailed native yeast protein is yet to be identified. Supporting a fundamental role of RQC in mitochondrial regulation, a yeast study suggested that mitochondrial mRNAs undergoing co-translational import are subjected to RQC regulation (Izawa et al., 2017).

Shared features between C-I30 MISTERMINATE in metazoans and CAT-tailing in yeast include dependency on RQC2 and regulation by VCP and Vms1/ANKZF1, non-templated elongation, preferential incorporation of A and T, and sensitivity to Anisomycin. Distinct features of C-I30-u MISTERMINATE include the use of AAs other than A and T, modification of a full-length protein, and induction by mitochondrial stress-induced eRF1/ABCE1 impairment. The mechanism underlying AA selectivity in CAT-tailing or MISTERMINATE is currently unknown.

In *PINK1* muscle, C-I30-u can be assembled into C-I, leading to impaired OxPhos. Moreover, it forms extra-mitochondrial aggregates and perturbs proteostasis. C-I30-u does not seem to be ubiquitinated. This is unlikely due to inhibition of ubiquitination by the CTE, as CAT-tailing can promote ubiquitination of cytosolic non-stop GFP in yeast (Kostova et al., 2017; Osuna et al., 2017). Lack of ubiquitination of C-I30 is likely due to inability of the RQC system to access ubiquitination sites, because C-I30 undergoes co-translational import such that its K residues are shielded from E3 ligases by the ribosomes and TOM/TIM. Other mechanisms, such as Vms1/ANKZF1-mediated cleavage of stalled peptidyl-tRNA (Verma et al., 2018), may play more important roles in MOM-associated RQC. Consistent with this notion, we show that ANKZF1 critically regulates C-I30-u formation and release from MOM, in a Parkin-regulated manner.

### Linking Mitochondrial Dysfunction with CTE-Induced Neurodegeneration

Our results indicate that once C-I30-u escapes the surveillance of RQC, it has the potential to form cytosolic aggregates. Such aggregates may attract other aggregation-prone proteins to form disease-specific aggregates. A “seeding” mechanism, in which a cascade of molecular events link mitochondrial dysfunction to proteostasis failure, mediated by CTE-initiated sequestration and aggregation of metastable proteins, may underlie CTE-induced neurodegeneration (Figure S7K). Supporting this model, mitochondrial proteins were found in disease-associated protein aggregates (Sergeant et al., 2003).

Our studies identify eRF1/ABCE1 as key factors transducing mitochondrial stress signals to the RQC machinery during MISTERMINATE. Previously, yeast eRF3 (Sup35) is shown to form prion aggregates [*PSI+*] (Glover et al., 1997), which sequesters WT eRF3 away from its normal function. However, the formation of [*PSI+*] is a rare event even under stress, and there is no indication that metazoan eRF3 can exist in a prion state. We did not find obvious alteration of eRF3 during MISTERMINATE. Instead, we observed reduced levels of eRF1 and ABCE1, and possible proteolytic cleavage of eRF1. ABCE1 is a Fe-S protein. Fe-S association is essential for ABCE1 function (Alhebshi et al., 2012), and the Fe-S domain of ABCE1 interacts with the C-term domain of eRF1 (Preis et al., 2014). ABCE1 may be part of the translation machinery that senses mitochondrial health. Indeed, metazoan ABCE1 is ubiquitinated under mitochondrial stress (Wu et al., 2018), and at least in yeast cells, WT level of ABCE1 is below the threshold sufficient for complete ribosome recycling (Young et al., 2015). ABCE1 is innately metastable and prone to sequestration by disease-associated aggregates (Olzscha et al., 2011). Sequestration of ABCE1 by CTE-induced aggregates may form a positive feedback loop that causes RQC failure. Our results indicate that Parkin plays an important role in regulating ABCE1/eRF1 complex formation, indicating a multifaceted role of PINK1/Parkin in MOM-associated translation.

### Role of Ribosome Release/Recycling Factors in Co-translational QC

Our results highlight the importance of termination/ribosome recycling in co-translational QC. A common step in various types of co-translational QC is splitting of stalled ribosomes before the clearance of NPCs attached to 60S (Schuller and Green, 2017). Whether a CTE process is involved in handling all 60S-associated NPCs remains to be seen. The eRFs are involved in resolving more than one type of stalled ribosomes. During NMD, SMG1 and UPF1 interact eRF1/eRF3 to form the SURF complex, which interacts with the exon junction complex (EJC) containing UPF2, UPF3, eIF4AIII, Y14, and Magoh, to trigger degradation of PTC-containing mRNAs. The faulty NPCs are also degraded during NMD (Kuroha et al., 2009), but the mechanism is poorly understood. Two EJC components, UPF3 and Y14, act as strong suppressors of C-I30-u formation and *PINK1* phenotypes when overexpressed. UPF3 knockdown is sufficient to induce C-I30-u and phenocopy *PINK1* mutant, which can be rescued by eRF1-OE. UPF3 interacts with eRFs (Neu-Yilik et al., 2017). Further studies of UPF3 will test whether its activity in termination is important for C-I30-u regulation.

Studies in yeast showed that the ribosome termination/recycling factor Dom34, the yeast homolog of Pelo, functions in rescuing ribosomes wondering past the stop codon, and that Dom34-ABCE1 interaction limits translation re-initiation in the 3′-UTR (Young et al., 2015). We show that Pelo-OE can reduce C-I30-u formation. It would be interesting to test if mitochondrial stress impacts 3′-UTR translation events by altering Pelo-ABCE1 interaction.

### Involvement of Co-translational QC in Disease

Our discovery of C-I30 MISTERMINATE was led by fortuitous observation of an aberrant form of C-I30 in *PINK1* flies, a PD model featuring prominent mitochondrial pathology. This emphasizes the importance of RQC to mitochondria biology and PD. PINK1 is extensively studied as a key regulator of mitochondrial QC, particularly in mitophagy (Pickrell and Youle, 2015). Our recent studies reveal that RQC and mitophagy are mechanistically linked, representing different steps in a continuum of organelle QC (Wu et al., 2018). Inefficient RQC may lead to accumulation of C-I30-u and possibly other CTE-carrying proteins such as ATP5a-u, which, if not timely removed from MOM, can result in activation of UPS/autophagy machineries and eventual mitophagy. A corollary of our finding is that neuronal degeneration caused by defective co-translational QC (Chu et al., 2009; Ishimura et al., 2014) may involve mitochondrial dysfunction. While MISTERMINATE may explain the intimate link between mitochondrial dysfunction and proteostasis failure, a fundamental question is whether the accumulation of CTE forms of many cellular proteins is needed, or that a selective few (e.g., C-I30-u and ATP5A-u) is sufficient, to cause neurodegeneration.

Due to the heavy burdens imposed on cellular QC systems by genomic alterations and increased protein synthesis in cancer cells, proteostasis is also highly relevant to cancer biology. Mitochondrial alterations are common in cancer cells (Deshaies, 2014), where MISTERMINATE may link mitochondria and protein homeostasis. Indeed, there is evidence implicating various RQC factors in cancer (Tian et al., 2016; Fessart et al., 2013). In the case of Rqc2/Clbn, we previously showed that it is induced by irradiation in a p53-dependent manner and regulates DNA damage-induced apoptosis (Wang et al., 2013b), and that it could suppress tumor growth in a rodent cancer model (Wang et al., 2013b).

Our results suggest that mitochondrial diseases, long thought to arise primarily from defective bioenergetics, could involve proteostasis failure as key disease mechanism; conversely, prevalent protein-aggregation diseases like AD/ALS could involve altered mitochondrial function. The RQC pathway may offer new therapeutic targets. In particular, Rqc2/Clbn/NEMF offers an attractive drug target given its essential role in the CTE process, the protective effect of its inactivation in a PD model as shown here, and its multi-domain structure potentially amenable to pharmacological intervention. Further investigation into the cause of eRF1/ABCE1 deficiency under stress may also lead to therapeutic strategies by boosting activities of eRF1/ABCE1 or other co-translational QC factors.

## ACKNOWLEDGEMENTS

We are grateful to Drs. Ramanujan Hegde and Susan Shao for providing plasmids and antibodies; Drs. Jiahuai Han and Elmar Wahle for antibodies; Drs. Yuzuru Imai, Ming Guo, and Richard Youle for cell lines; Drs. Jongkeong Chung, William Saxton, Serge Birman, Patrick Verstreken, Rongwen Xi, GW 2^nd^ Dorn, Mika Rämet, Paul Taylor, the Vienna *Drosophila* RNAi Center, FlyORF, and the Bloomington *Drosophila* Stock Center for fly stocks; Dr. Ryan Lieb and Kratika Singhal from Stanford University Mass Spectrometry for Mass Spec. Dr. Lihua Jiang for identifying the CID peptide spectra. Special thanks go to J. Gaunce for maintaining flies and providing technical supports and members of the Lu lab for discussions. Supported by the NIH (R01NS084412 to B.L. and R01GM115968 to O.B.).

## AUTHOR CONTRIBUTIONS

Z.W designed the study, performed the experiments, analyzed data, and co-wrote the manuscript; J.L, I.T, Y.L, J.D, Z.D, C.S performed experiments and analyzed data; S.C, O.B analyzed data; M.S and O.B provided resources for analyzing data; X.B, S.G., G.A. provided key reagents; B.L conceived and supervised the study, performed experiments, wrote the manuscript, and provided funding.

Correspondence and requests for materials should be addressed to B.L.

## EXPERIMENTAL PROCEDURES

### Fly Stocks and Fly Culture condition

*dPINK1*^*B9*^ mutant was a gift from Dr. Jongkeong Chung. *dparkin*^*1*^ and *dparkin*^*Δ 21*^ mutants were from Dr. Patrik Verstreken. *TH-Gal4* was a gift from Dr. Serge Birman. *UAS-mito-GFP* was from Dr. William Saxton. *UAS-dPINK1 RNAi* was generated in our lab (Gehrke et al., 2015). *UAS-dVCP WT, dVCP RNAi* and *UAS-dVCP DN* were gifts from Dr. Paul Taylor. *UASp-Pelo* was kindly provided by Dr. Rongwen Xi. *UAS-dNOT4-myc (M)* and *UAS-NOT4 (W)* were from Dr. Mika Rämet. *clbn+/-* and *UAS-clbn* flies were described before (Wang et al., 2013b). *UAS-hMfn1* was a gift from Dr. GW 2nd Dorn. UAS-*dDrp1 DN* was generated in our lab.

*dABCE1-HA* (F001097) and *dC-I30-HA* (F003000) were obtained from FlyORF. *dPelo RNAi* (v108606), *dNOT4 RNAi* (v110472), *dAARS RNAi* (v17171), *dTARS RNAi* (v7752), *dGARS RNAi* (v106748), *dHARS RNAi* (v104672), *dMARS RNAi* (v106493), *dVARS RNAi* (v21782), *dQARS RNAi* (v101761), *dP/EARS RNAi* (v34995), *dLARS RNAi* (v45048), *dSARS RNAi* (v41928), *dYARS RNAi* (v105615), *dRARS RNAi* (v42185), *dFARS RNAi* (v107079), *dWARS RNAi* (v107049), *dCARS RNAi* (v45611), *dND-SGDH RNAi* (v11381, v101385), *dATPsynC RNAi* (#57705), *blw RNAi* (v34663) and *deRF3 RNAi* (v106240) were from the VDRC. dC-I30 RNAi (#44535, #51425), *dNOT4 EP* (#22246), *dABCE1 RNAi* (#31601), *dY14 EP* (#6597), *dY14 RNAi* (#55367), *deRF1 EP* (#17265), *deRF1 RNAi* (#67900), *deRF1*^*KY7*^ mutant (#7069), *deRF3 EP* (#20740), *deRF3*^*LR17*^ mutant (#7067), *UAS-GFP-dUPF1 (#24623), dUPF1 RNAi* (#64519), *dUPF3 EP* (#16558), *dUPF3 RNAi* (#44565), *dXRN1 EP* (#33263), *dXRN1 RNAi* (#34690), *dATPsynO RNAi* (#43265) and *dKARS RNAi* (#32967) and other general fly stocks were from the Bloomington *Drosophila* Stock Center.

Fly culture and crosses were performed according to standard procedures and raised at indicated temperatures.

### Fly Behavior Test and ATP Measurement

For all wing posture assays, male flies at 7-day and 14–15-day old and aged at 29°C were used. 20 male flies were collected/raised in one vial and 3 independent vials were counted *per* genotype. Measurements of ATP contents in thoracic muscle were performed as described previously (Gehrke et al., 2015), using a luciferase based bioluminescence assay (ATP Bioluminescence Assay Kit HS II, Roche Applied Science). 3 thoraces were used for one assay and 3∼4 assays were performed for each genotype.

### Fly Muscle and Brain Staining

For immunohistochemical analysis of mitochondrial morphology of adult fly brains and muscle tissues, 5-day old male flies from 29°C were assayed. In muscle staining, at least 5 individuals were examined for each genotype and the representative images were presented. For analysis of DA neuron number in the various genetic backgrounds, 21-day old male flies were assayed. In DA neuron staining, at least 7 individuals were examined for each genotype. Dissected tissue samples were briefly washed with 1x PBS and fixed with 4% formaldehyde in 1x PBS containing 0.25% Triton X-100 for 30 minutes at room temperature. Fixatives were subsequently blocked with 1x PBS containing 5% normal goat serum and incubated for 1 hour at room temperature followed by incubation with primary antibodies at 4°C overnight. The primary antibodies used were: chicken anti-GFP (1:5,000, Abcam), rat anti-HA (1:1,000; Sigma), rabbit anti-TH (1:1000, Pel-Freez), and mouse anti-C-I30 (1:1000, Abcam). After three washing steps with 1x PBS/0.25 % Triton X-100 each for 15 minutes at room temperature, the samples were incubated with Alexa Fluor® 594-conjugated, Alexa Fluor® 488-conjugated, and Alexa Fluor® 633-conjugated secondary antibodies (1:500, Molecular Probes) for 3 hours at room temperature and subsequently mounted in SlowFade Gold (Invitrogen).

### Cell Lines, Cell Culture and Cell Transfection Condition

Regular HeLa cells and *PINK1*^-/-^ HeLa cells (a gift from Dr. Richard Youle, NINDS) were cultured under normal conditions (1x DMEM medium, 8.75% FBS, 37°C, 5% CO_2_). HeLa/GFP-Parkin cells (a gift from Dr. Yuzuru Imai, Juntendo University) were cultured in the same condition with additional drug selection (1µg/ml puromycin).

Cell transfections were performed by using Lipofectamine 3000 reagent (cat#: L3000015, Invitrogen) and knocking-down experiments were performed using Lipofectamine RNAiMAX reagent (cat#: 13778150, Invitrogen), according to instructions from the manufacturers. Stealth RNAi™ siRNAs (Invitrogen) used for the RNAi experiments are: CNOT4 (HSS107246, HSS107245), ABCE1 (HSS109286, HSS109285), siRNAs: siCON (cat#:12935-400, Invitrogen), siNEMF (NEMFHSS113541, NEMFHSS113540), siVCP (VCPHSS111263, VCPHSS111264), siHARS (HARSHSS104694), siAARS (AARSHSS100022), siTARS (TARSHSS110482), siYRAS (YARSHSS112540), siSARS (SARSHSS109468) from Invitrogen Inc.

### Immunohistochemical Analysis of cultured cells

For immunohistochemical analysis in human cells, the cells were cultured on the ethanol-cleaned cover glass. Cells were washed with 1xPBS and fixed with 4% formaldehyde in 1x PBS for 30 minutes at room temperature, later washed and permeabilized with 1x PBS containing 0.25% Triton X-100 for 15 minutes. The fixed samples were subsequently blocked with 1x PBS containing 5% normal goat serum and incubated for 1 hour at room temperature followed by incubation with primary antibodies at 4°C overnight. The primary antibodies used were mouse anti-Flag (1:1,000, Sigma-Aldrich), rabbit anti-Flag (1:1,000, Sigma-Aldrich), rabbit anti-TOM20 (1:1,000, Santa Cruz), rabbit anti-BTF3 (1:500, Abcam), guinea pig anti-P62 (1:500, ProGEN), mouse anti-RpS6 (1:500, Cell Signaling), and rabbit anti-RpL3 (1:500, Cell Signaling). The secondary antibodies used were Alexa Fluor® 488, 594 and 633-conjugated antibodies (1:500, Molecular Probes).

### Drug Treatments

For drug treatment of *Drosophila*, flies were collected after eclosion and divided into separate vials (20∼25 flies each vial). Instant fly food (Carolina) was mixed with 1% H_2_O_2_ (cat#: H1009, Sigma-Aldrich), 250 µM rotenone (cat#: R8875, Sigma-Aldrich) or 100 µM CCCP (cat#: C2759, Sigma-Aldrich). Vials were changed every day. Samples were collected for further analyses after 5 days of treatment. In order to achieve high efficiency, anisomycin stock (100mM in DMSO, cat#: A9789, Sigma-Aldrich) were diluted with the Schneider’s Medium (cat#: 21720-024, Gibco™) to 250 μM and injected into fly thorax between mesonotum and scutellum *via* a hand-made glass needle. After injection, flies were kept in the vial with standard fly food and waited for 24 hours before the experiments.

For drug treatment in human cells, HeLa cells or HeLa/GFP-Parkin cells were cultured and treated with CCCP at the indicated concentrations and durations as described (Narendra et al., 2008). For most experiments, 20 µM CCCP concentration was used. For Antimycin/Oligomycin treatment, 10 µM of each compound was used. For anisomycin, 200 µM was used, and for the VCP inhibitor NMS-873 (cat#: S7285, Selleckchem), 1 µM was used.

### SDS-PAGE Sample Preparing, Gel Preparing and Running Conditions

Home-made gels: The home-made Laemmli resolving gel (10 ml) contains 3 ml of 40% Acrylamide/Bis (cat#: 1610148, Bio-Rad), 2.5 ml of 1.5M Tris-HCl pH 8.8 (cat#: 1610798, Bio-Rad), 1 ml of 10% SDS (cat#: 1610416, Bio-Rad), 50 µl of 10% APS (cat#: 1610700, Bio-Rad) and 5 µl of TEMED (cat#: 1610800, Bio-Rad). The stacking gel (2.5 ml) contains 0.25 ml of 40% Acrylamide/Bis, 0.63 ml of 0.5M Tris-HCl pH 6.8 (cat#: 1610799, Bio-Rad), 250 µl of 10% SDS, 12.5 µl of 10% APS and 2.5 µl of TEMED. The running buffer (1L) contains 3g Tris base (cat#: 11814273001, Sigma-Aldrich), 14.4g Glycine (cat#: G8898, Sigma-Aldrich), and 1g SDS (cat#: L3771, Sigma-Aldrich).

Commercial gels: NuPAGE™ 4-12% Bis-Tris Protein Gels (cat#: NP0321BOX, Invitrogen) and NuPAGE® MOPS SDS running buffer (cat#: NP0001, Invitrogen) was used for SDS-PAGE and immunoblot analyses. NativePAGE™ 3-12% Bis-Tris Protein Gels (cat#: BN1001BOX, Invitrogen) and NativePAGE™ Running Buffer Kit (cat#: BN2007, Invitrogen) were used for Native-PAGE as the first dimension of the 2-D gel. The gel slices were cut and denatured by heating in the 1x Laemmli SDS sample buffer for 10 minutes and then load to Home-made SDS-PAGE for the second dimension.

### Western Blots and Antibodies

The following antibodies were used for western blot analyses: anti-Actin (cat#: A2066, Sigma-Aldrich), anti-Tubulin (cat#: MA1-80017, Thermo Fisher), anti-CI-30 (cat#: ab14711, Abcam), anti-FLG (cat#: F1804, F7425, Sigma-Aldrich), anti-HA (cat#: 11867423001, Sigma-Aldrich), anti-GFP (cat:# ab13970, Abcam), anti-TH (cat#: P40101-150, Pel-Freeze), anti-eRF1 (cat#: 13916S, Cell Signaling), anti-eRF3 (cat#: 14980S, Cell Signaling), anti-ABCE1 (a gift from Dr. R Hegde), anti-Pelo (cat#: ab140615, Abcam), anti-Hsp90 (cat#: A5015, Selleckchem), anti-Hsp60 (cat#: sc-59567, Santa Cruz), anti-Hsp70 (cat#: A5002, Selleckchem), anti-BTF3 (cat#: ab66940, Abcam), anti-P62 (cat#: GP62-C, ProGEN), anti-eIF4G (cat#: 2498S, Cell Signaling), anti-RpS6 (cat#: 2317S, Cell Signaling), anti-RpL3 (cat#: sc-86828, Santa Cruz Biotech), anti-TOM20 (cat#: sc-17764, sc-11415, Santa Cruz Biotech), anti-ATP5a (cat#: ab14748, Abcam); anti-SDHA (cat#: 14715, Abcam); anti-NDUFV1 (cat#: NBP1-33074, Novus); anti-pS51-eIF2α (cat# S9721, Cell Signaling); anti-eIF2α (cat# 9722, Cell Signaling); anti-pS127-YAP (cat# 13008T, Cell Signaling); anti-ANKZF1 (cat# sc-398713, Santa Cruz), anti-Clbn (a gift from Dr. Bi).

To obtain quantitative western blot results, experimental conditions such as the amounts of protein loaded, the antibody dilutions, and exposure times were adjusted to make sure that the western blot signals were in the linear range. For western blots, analyses were performed using standard protocols. After incubation with the ECL reagent (cat# NEL103E001EA, PerkinElmer Inc.), the signals were recorded on X-ray film (cat# E3018, Denville Scientific Inc.), scanned, and signal intensity analyzed by ImageJ. For all the blots used in the manuscript, at least three biological repeats were assayed, and the typical ones were presented. The numbers under the blots represent the ratio of presented blot. The statistical analyses in Figure 2J, 3G and S2E were based on data from three independent biological repeats.

### Co-IP Experiments

For C-I30-u immunoprecipitation, we transiently expressed *pcDNA3.1(+)-C-I 30-FLG-TEV* plasmid in HeLa cells. Sixty hours post-transfection, we applied UV cross-linking and 0.5% formaldehyde in 1x PBS to the attached cells on the petri dish. We homogenized the cells in the lysis buffer [50 mM Tris-HCl, pH7.4, 150 mM NaCl, 5 mM EDTA, 10% glycerol, 1% Triton X-100, 0.1mg/ml cycloheximide, 1x RNase inhibitor, and Complete protease inhibitor cocktail (cat#: B14011, Bimake)], additional Phosphatase Inhibitor Cocktail (cat#: B15001, Bimake) will be applied if phosphorylation signal is to be detected. After centrifugation at 10,000g for 5 min, the supernatant was subjected to immunoprecipitation using the indicated antibodies, or affinity gels (Anti-FLAG M2 affinity gel, cat#: A2220, Sigma-Aldrich) at 4°C for 6 hours with gentle shaking. Subsequently, the sepharose beads were washed three times (10 minutes each) at 4°C in lysis buffer, mixed with 2x SDS Sample buffer, and loaded onto SDS-PAGE gels.

### Mitochondria and CI-30-u Purification and C-I activity measurement

Intact mitochondria from *in vitro* cell cultures and fly muscle tissue samples were purified and quality controlled for the absence of contamination by other organelles as described previously (Gehrke et al., 2015). For analysis of fly samples, male flies at appropriate ages were used for thoracic muscle dissection. To block the release of mRNAs associated with mitochondria, 0.1 mg/ml cycloheximide was applied to all buffer solutions. Samples were homogenized using a Dounce homogenizer. After two steps of centrifugation (1,500g and 13,000g), the mitochondria pellet was washed twice with HBS buffer (5 mM HEPES, 70 mM sucrose, 210 mM mannitol, 1 mM EGTA, 1x protease inhibitor cocktail), then resuspended and loaded onto Percoll gradients. The fraction between the 22% and 50% Percoll gradients containing intact mitochondria were carefully transferred into a new reaction tube, mixed with 1 volume of HBS buffer, and centrifuged again at 20,000g for 20 minutes at 4°C to collect the samples for further analyses. The mitochondria then was solubilized by 5% Digtonin (cat#: BN2006, Invitrogen) on ice for 30 min and prepared as the sample for Blue Native PAGE using the NativePAGE™ Sample PreP Kit (cat#: BN2008, Invitrogen), or solubilized for C-I30-u purification.

For denature immunoprecipitation of fly C-I30-u, partially purified mitochondria sample was first extracted by mitochondrial lysis buffer (50 mM Tris-HCl, pH7.4, 150 mM NaCl, 5 mM EDTA, 10% glycerol, 1% NP-40, 0.1mg/ml cycloheximide, 1x RNase inhibitor, and Complete protease inhibitor cocktail) on ice for 15 minutes, and the remaining pellets were prepared by solubilization with 1x SDS Sample buffer. The samples were denatured by boiling at 100°C, and then diluted with pre-chilled lysis buffer at a 1:5 ratio and subjected to immunoprecipitation using anti-CI-30 (cat#: ab14711, Abcam).

Complex-I activity was measured using the Complex I Enzyme Activity Microplate Assay Kit (cat# ab109721, Abcam) and following the manufacturer’s protocol. 200µg of purified mitochondrial sample was used in each measurement. The colorimetric signals were read using the Epoch plate reader (BioTek, USA).

For mitochondrial ribosomal fraction purification, briefly, purified mitochondria from human cells were lysed with Buffer A (20 mM HEPES, 50 mM KCl and 10 mM MgCl_2_) with additional 0.1% Triton X-100, 0.1 mg/ml cycloheximide and 1x RNase Inhibitor. The mitochondrial lysate was pre-cleared by centrifugation (20,000g, 10 min, 4°C), and then loaded onto a 25% sucrose cushion (made in Buffer A). The ribosome fraction was pelleted via high-speed centrifugation (70,000 rpm, TLA 100.3 Rotor, 20 min, 4°C). After centrifugation, the ribosomal pellet was resuspended in Buffer A containing 1% Triton X-100 and analyzed by immunoblotting.

### Trypsin Partial Digestion, Mass Spec, and Database Searching Conditions

C-I30 IP samples were combined and run on a 4-12% SDS-PAGE gel and resolved protein bands stained by Coomassie blue R-250 staining. Protein bands were excised out into a 1.5 ml Eppendorf tubes and then cut into 1×1 mm squares. The excised gel pieces were then reduced with 5 mM DTT, 50 mM ammonium bicarbonate at 55°C for 30 min. Residual solvent was removed and alkylation was performed using 10 mM acrylamide in 50 mM ammonium bicarbonate for 30 min at room temperature. The gel pieces were rinsed 2 times with 50% acetonitrile, 50 mM ammonium bicarbonate and placed in a speed vac for 5 min. Digestion was performed with Trypsin/LysC (Promega) in the presence of 0.02% protease max (Promega) in both a standard overnight digest at 37°C as well as in a limited digest format (2 hours at 50°C). Samples from both digestion conditions were combined to one tube, centrifuged and the solvent including peptides were collected and further peptide extraction was performed by the addition of 60% acetonitrile, 39.9% water, 0.1% formic acid and incubation for 10-15 min. The peptide pools were dried in a speed vac. Samples were reconstituted in 12.5µl reconstitution buffer (2% acetonitrile with 0.1% Formic acid) and 3µl (100ng) of it was injected into the instrument.

LC/MS data of the trypsin-Partially-digested peptides were collected on an Orbitrap Fusion mass spectrometer (Thermo Scientific, San Jose, CA) with a Nanoacquity M Class UPLC (Waters Corporation, Milford, MA). For a typical experiment, a flow rate of either 300 nL/min or 450 nL/min was used, where mobile phase A was 0.2% formic acid in water and mobile phase B was 0.2% formic acid in acetonitrile. Analytical columns were prepared in-house with an I.D. of 100 microns pulled to a nanospray emitter using a P2000 laser puller (Sutter Instrument, Novato, CA). The column was packed using C18 reprosil Pur 2.4 µM (Dr. Maisch) to a length of ∼25 cm. Peptides were directly injected onto the analytical column using a linear gradient (4-40% B) of 80min. The mass spectrometer was operated in a data dependent fashion using CID fragmentation for MS/MS spectra generation collected in the ion trap with a collisional energy of 35%.

### Customized Database Searches

#### For the Generic Database Searches

In a typical LCMS data analysis, the .RAW data files were processed using Byonic v 2.14.27 (Protein Metrics, San Carlos, CA) to identify peptides and infer proteins. Proteolysis was assumed to be semi-tryptic with up to four missed cleavage sites. Precursor mass accuracies were held within 12 ppm, and 0.4 Da for MS/MS fragments. Proteins were held to a false discovery rate of 1%, using standard approaches (Elias and Gygi, 2007). Resulting peptides were then investigated for potential modification using either wildcard search conditions, or modified .fasta files including potential peptide sequences of interest (see text).

#### For the Customized Database Searches

Raw data were analyzed using the Thermo Proteome Discoverer computational software (version 1.4.1.14) with Sequest HT as search engine. A precursor mass tolerance was set to ±12 ppm and a fragment mass tolerance of ±0.8 Da. We allowed up to 4 missed tryptic cleavages for trypsin semi-digestion. The maximum false peptide discovery rate was specified as 1% (FDR<0.01). The resulting peptide files were searched against in-house peptide sequencing pools, based on our finding of ARS-dependency of C-I30-u formation. Identified peptides were filtered for high confidence, a search engine rank of 1 and peptide mass derivation of 7 ppm. All the peptide assignments with interest were also manually validated. The proteomics data can be accessed via ProteomeXchange (PXD*). The source code (requiring Python environment) to generate the in-house peptide sequencing pools will be available upon request.

### Plasmids and Molecular Cloning

The original human C-I30 CDS sequence was from the *pPM-N-D-C-His* (PV394217, abm) plasmid. *pcDNA3.1(+)-C-I30-FLG* and *pcDNA3.1(+)-C-I30-FLG-TEV* plasmids were generated by cloning the human C-I30 CDS with different tags into *pcDNA3.1(+)* vector via Kpn I and Xba I sites.

*pcDNA3.1(+)-C-I30-TEV-FLG*; *pcDNA3.1(+)-C-I30-FLG-AT*_*5*_; *pcDNA3.1(+)-C-I30-FLG-AT*_*23*_; *pcDNA3.1(+)-C-I30-FLG-non-AT*_*25*_; *pcDNA3.1(+)-C-I30(mMTS)-FLG (36R*→*A)*; *pcDNA3.1(+)-C-I30-FLG-A*^*stop*^; *pcDNA3.1(+)-C-I30-FLG-AK*^*stop*^; *pcDNA3.1(+)-C-I30-FLG-AKK*^*stop*^ and *pcDNA3.1(+)-HA-C-I30-FLG*; were modified based on the *pcDNA3.1(+)-C-I30-FLG-TEV* plasmid *via* the Q5 Site-Directed Mutagenesis Kit (cat#: E0554S, NEB). *pCMV-FLAG-ABCE1* was a gift from Dr. Ramanujan Hegde. *pCMV6-FLAG-NOT4* and *pCMV6-ANKZF1* was obtained from OriGene Inc (cat#: RC217418 and RC201054 TrueORF). *pCMV-SPORT6.1-eRF1* (cat#: MHS6278-202804766, Dharmacon™) and *pCMV-SPORT6.1-VCP* (cat#: MHS6278-202760239, Dharmacon™) were from GE healthcare. All plasmids have been confirmed by sequencing.

### CLIP assay and tRNA RT-PCR

CLIP assays were performed as we described before (Gehrke et al., 2015). Briefly, we placed the fly lysates on ice and performed UV cross-linking using a Stratalinker 2000. We then performed regular IP, washed the beads and added 2xSDS buffer to the beads as usual. Samples were loaded onto 12% SDS-PAGE gel and prepared for WB. After WB, we cut out a small section from the PVDF membrane covering the expected protein position of Clbn. We prepared 4mg/ml proteinase K in 1xPBS and add 30ul to the cut out membrane, incubated 20min at 37°C. We then extracted RNA from the membrane strip by following manufacturer’s manual of the RNA extraction kit (RNeasy kit, cat# 74104, Qiagen) by adding 100ul of RLT+ β-mercapto-ethanol. We used a homogenizer to enhance RNA removal from the membrane. We then used the purified RNA samples for RT-PCR with the OneStep RT-PCR kit (cat# 210210, Qiagen).

The following primers were used in RT-PCR analyses to detect fly tRNAs from the CLIP assay: RT-tRNA(Ala_AGC)-N: *GGGGATGTAGCTCAGATG*; RT-tRNA(Ala_AGC)-C: *TGGAGATGCGGGGTATCG*; RT-tRNA(Ala_CGC)-N: *GGGGACGTAGCTCAGTG*; RT-tRNA(Ala_CGC)-C: *TGGAGACGCCGGGGTTT*; RT-tRNA(Ala_TGC)-N: *GGGGATGTAGCTCAGTGG*; RT-tRNA(Ala_TGC)-C: *TGGAGATGCCGGGGATCG*; RT-tRNA(Thr_AGT)-N: *GGCGCCGTGGCTTAGTT*; RT-tRNA(Thr_AGT)-C: *AGGCCCCGGCGAGATTC*; RT-tRNA(Thr_CGT)-N: *GCCTCTTTAGCTCAGTGG*; RT-tRNA(Thr_CGT)-C: *TGCCTCCTGTGAGGGTTG*; RT-tRNA(Thr_TGT)-N: *GCCTCTTTAGCTCAGTGG*; RT-tRNA(Thr_TGT)-C: *TGCCTCCTGTGAGGATTG*; RT-tRNA(Ser_CGA)-N: *GCAGTCGTGGCCGAGTG*; RT-tRNA(Ser_CGA)-C: *CGCAGCCGGTAGGATTCG*; RT-tRNA(Ser_AGA)-N: *GCAGTCGTGGCCGAGCG*; RT- tRNA(Ser_AGA)-C: *CGCAGTCGGTAGGATTCG*; RT-tRNA(Ser_TGA)-N: *GCTGCGGTGTCCGAGTG*; RT-tRNA(Ser_TGA)-C: *CGCTGCGGACAGGACTTG*.

### PPTase, USP2 treatment, Enterokinase digestion and TEV Digestion

For Phosphatase reaction, the purified C-I30-Flag protein (30µl beads) was digested on beads (Anti-FLAG M2 affinity gel) with final 10U CIP Phosphatase (cat#: M0290S, NEB) in the 1x CIP buffer (50 mM Potassium Acetate, 20 mM Tris-acetate, 10 mM Magnesium Acetate, 100 ug/ml BSA, pH 7.9 at 25°C). The reaction was performed for 30 minutes at 37°C with gentle shaking.

For USP2 reaction, the purified C-I30-Flag protein (30µl beads) was pre-equilibrated with 1X DUB reaction buffer (50 mM Tris pH 7.5, 50 mM NaCl and 5 mM DTT) and digested on beads (Anti-FLAG M2 affinity gel). USP2 was diluted in 50 mM Tris pH 7.5, 300 mM NaCl and 2 mM DTT buffer and pre-incubated for 10 min at 22C. Subsequently, each sample was incubated with pre-incubated 3 µM USP2 (cat#: E-504, BostonBiochem) for different time points (0.5, 1 and 2 hours) at 37°C with gentle shaking.

For Enterokinase reaction, the purified C-I30-Flag protein (30µl beads) was digested on beads (Anti-FLAG M2 affinity gel) with final 5U Enterokinase (cat#: P8070S, NEB) in the 1x EK reaction buffer (20 mM Tris-HCl, 50 mM NaCl, 2 mM CaCl_2_ pH 8.0). The reaction was performed for 16 hours to balance the efficiency and specificity at 22°C with gentle shaking.

For TEV digestion, the purified C-I30-Flag protein (30µl beads) was digested on beads (Anti-FLAG M2 affinity gel) with 10U ProTEV Plus (cat#: V6101, Promega) in the 1x ProTEV digestion buffer (50 mM HEPES pH 7.0, 0.5 mM EDTA). The reaction was performed for 6 hours to achieve the maximal efficiency at 30°C with gentle shaking.

After reaction, the supernatant was separated from the beads *via* centrifugation. Beads were washed once with 1x PBS. The samples were mixed with 2x SDS sample buffer and denatured by heating, and then loaded for western blot analyses.

### Quantification and Statistical Analysis

All analyses were performed with SPSS (IBM, USA) and further confirmed by MATLAB (MathWorks, USA). Error bars represent standard deviation (SD). Data in Figure 5D, 5K and Supplemental Figure S5H, S7D and S7F were analyzed by *Chi-squared* test. For pair-wise comparisons in Figure 3E, S3A, S4C, S4E we used two-tailed Student’s *t*-test. For comparing multiple groups as in Figure S1C, 2E, 2H, 2J, S2F, 3G, S4E, 5B, S5C and S5D we used one-way ANOVA test followed by Student–Newman–Keuls test (or SNK test) plus Bonferroni correction (multiple hypotheses correction). In our statistical comparisons, * indicating *p* < 0.05 and ** indicating *p* < 0.01 were considered as significant differences. Sample size/statistical details were stated separately in the figure legends.

