## supplemental figures and legend for "MISTERMINATE Mechanistically Links Mitochondrial Dysfunction with Proteostasis Failure"

### SUPPLEMENTAL FIGURE LEGEND

**Figure S1. Effects of Knocking down ATP Synthetase Subunits on C-I30-u Expression and Genetic Manipulation of RQC Genes in Muscle Tissues**

(A) Immunoblots of thoracic muscle samples from flies with OxPhos-related genes knocked down by RNAi in *PINK1* mutant *(dPINK1-/Y)* background. Actin serves as loading control.

(B) Immunostaining of fly muscle tissue showing mitochondrial morphology changes by the manipulation of co-translational quality control genes in wild type and *Mhc-Gal4>PINK1* RNAi conditions. Mitochondrial morphology is monitored with a mito-GFP reporter. Scale bar, 5μm.

(C) Induction of abnormal wing posture by manipulating select co-translational quality control factors in wildtype background. * or #, $p<0.05$ in SNK-test plus Bonferroni correction vs. *control (w-)* group at day 7 or day 15, respectively.

**Figure S2. Further Evidence Supporting Translational Control of C-I30-u Formation**

(A) Immunoblots of HA (C-I30-HA) and C-I30 in wild type or *PINK1* mutant backgrounds showing the formation of C-I30-u for both endogenous and exogenous C-I30 proteins. Actin serves as loading control. (short) and (long) stand for shorter and longer exposures.

(B) Immunoblots of Flag (C-I30) in C-I30-Flag transfected HeLa cell showing the time-dependent accumulation of C-I30-u upon CCCP or antimycin A/oligomycin (A/O) treatment. Tubulin serves as loading control.

(C) Immunoblots of C-I30-3xFlag or C-I30(mMTS)-3xFlag in vehicle-, CCCP-, or A/O-treated HeLa cells. C-I30(mMTS)-3xFlag has the R36 residue (underlined below) in the MTS cleavage site mutated to A (R36A):

MAAAAVARLWWRGILGASALTRGTGRPSVLLLPVRRESAGADTRPTVRPRNDVAHKQLSA

(D) Immunoblots showing elevated protein expression of the RQC genes (eRF1, and UPF1) in the overexpression fly lines.

(E) Immunoblots of muscle samples from *Mhc-Gal4* driven RNAi transgenes of *ABCE1*, *Pelo*, *NOT4*, *Y14*, *UPF1*, *UPF3*, *eRF1*, *eRF3*, *XRN1* and *VCP* in *PINK1* mutant background. *w*- serves as wildtype control and Actin as loading control.

(F) Quantification of data shown in E.

**Figure S3. Further Characterization of the Nature of CTE in C-I30-u**

(A) Restoration of ATP level by *clbn* mutation in *PINK1* mutant background. * $p<0.05$; Student's *t*-test. Values are normalized to *clbn (-/+)* heterozygous control.

(B) Rescue of DA neuron mitochondrial morphology by knocking down *clbn* in *PINK1* mutant background. Scale bar, 3μm.

(C) Immunostaining of fly muscle tissue showing the effect on mitochondrial morphology by inhibiting various tRNA synthetases in *Mhc-Gal4>PINK1* RNAi animals. Scale bar, 5μm.

(D) Immunoblots showing the removal of C-I30-u by anisomycin treatment of *hMfn1 OE*, *UPF3 RNAi* and *VCP DN* transgenic flies driven by *Mhc-Gal4*.

(E) Immunoblots of HeLa cells transfected with Flag-tagged C-I30-AK$^{stop}$, -A$^{stop}$, or -AKK$^{stop}$ and treated with or without CCCP.

(F) Diagram showing the C-I30-TEV-Flag and C-I30-Flag-TEV constructs, and immunoblots of HeLa cell lysates transfected with each plasmid showing the induction of C-I30-u upon CCCP treatment. Actin serves loading control.

(G) Immunoblots of Flag IP samples from HeLa cell transfected with C-I30-TEV-Flag or C-I30-Flag-TEV and treated with or without TEV protease, showing the removal of C-I30-u in C-I30-

Flag-TEV sample, and release of two Flag-positive fragments into the supernatant (sn) from digested CI-30-TEV-Flag sample. Samples were treated with formic acid to dissociate aggregates. Blue *: Flag-CTE; blue >: Flag only. These data further support the CTE model of C-I30-u formation. Note that due to its small size, some of the Flag only fragment might have been lost during transfer and processing.

(H) Immunoblots of detergent soluble and insoluble fractions from *PINK1* whole fly extracts showing enrichment of C-I30-u in the detergent insoluble fraction. Commassie blue R-250 staining shows C-I30-u protein purified by denaturing IP and used for Mass Spec.

(I) The MS/MS spectrum of the parent ion of peptides, showing b ions and y ions labeled in the spectrum. Based on MS/MS sequencing, the matched entries in the customized database were shown on the upper left.

In the MS/MS spectrum of 899.99670 Da ion ($MH^+$ = 1798.98613 Da; putative ID: NANPPAEVVPPQAPAKKA), mass/charge ratios of observed b4, b5, b8-10 ions were 397.18, 494.24, 793.38, 892.45, 989.51 Da, respectively, those of observed $b8^{2+}$, $b10^{2+}$, $b11^{2+}$, $b13^{2+}$, $b17^{2+}$ ions were 397.20, 495.26,543,78, 643.33,723.38, 855.47 Da. Mass/charge ratios of observed y6, y9-11, y13–15 ions were 585.37, 907.54, 1006.60, 1105.67, 1305.75, 1402.81, 1499.86 Da, respectively, those of observed $y15^{2+}$, $y16^{2+}$, $y17^{2+}$ ions were 750.43, 807.45, 842.97 Da.

In the MS/MS spectrum of 721.37537 Da ion ($MH^+$ = 2162.11155 Da; putative ID: PAEVVPPQAPAKKAAAEETEE), mass/charge ratios of observed b4-9 ions were 397.20, 496.28, 593.33, 690.38, 818.44, 899.48 Da, respectively, those of observed $b9\text{-}14^{2+}$, $b16^{2+}$, $b17^{2+}$, $b20^{2+}$ ions were 445.24, 493.77, 529.29, 593.33, 657.38, 692.90, 763.94, 828.46, 1008.03 Da. Observed $b8^{3+}$, $b13^{3+}$, $b15\text{-}17^{3+}$, $b19^{3+}$ ions were 273.49, 438,59, 485.95, 509.63, 552.64,

629.34 Da. Mass/charge ratios of observed y4-8, y10 ions were 507.19, 636.24, 707.27, 778.31, 849,35, 1105.54 Da, respectively, those of observed y10-12$^{2+}$, y14$^{2+}$, y15$^{2+}$ y17$^{2+}$, y18$^{2+}$, y20$^{2+}$ ions were 553.27, 588.79, 637.32, 736.87, 785.39, 883.45, 932.99, 1033.03 Da. Observed y9$^{3+}$, y11$^{3+}$, y13$^{3+}$, y16$^{3+}$, y17$^{3+}$, y19$^{3+}$ ions were 326.49, 392.86, 448.89, 556.28, 589.30, 665.34 Da.

In the MS/MS spectrum of 649.73096 Da ion (MH$^{+}$ = 3244.62568 Da; putative ID: KAAAAAAAAAAAAAAAAATTTTTTTTTTTTTTTT), mass/charge ratios of observed b5, b7-9 b12 ions were 413.25, 555.32, 626.36, 697.40, 910.51 Da, respectively, those of observed b8$^{2+}$, b10$^{2+}$, b13$^{2+}$, b17-19$^{2+}$ b21-23$^{2+}$ ions were 313.68, 384.72, 491.28, 633.35, 668.87, 704.39, 805.44, 855.96, 906.48 Da. Observed b12-14$^{3+}$, b16$^{3+}$, b18-28$^{3+}$ ions were 304.18, 327.85, 351.53, 398.89, 446.25, 469.93, 503.61, 537.29, 570.98, 604.66, 638.34, 672.02, 705. 71, 739.39, 773.07 Da. Observed b23$^{4+}$, b24$^{4+}$, b27$^{4+}$, b28$^{4+}$ ions were 453.75, 479.01, 554.79, 580.06 Da. Observed b15$^{5+}$, b26$^{5+}$, b28$^{5+}$ ions were 225.53, 423.83, 464.25 Da. Mass/charge ratios of observed y10-12$^{2+}$, y14$^{2+}$ ions were 515.25, 565.77, 616.30, 717.35 Da, respectively, those of observed y10$^{3+}$, y14$^{3+}$, y16-19$^{3+}$, y22$^{3+}$, y23$^{3+}$, y25-30$^{3+}$ ions were 343.84, 478.57, 545.93, 579.61, 613.30, 636.98, 708.01, 731.69, 779.05, 802.73, 826.41, 850.09, 873.77, 897.46 Da. Observed y11$^{4+}$, y14-16$^{4+}$, y18-20$^{4+}$, y23$^{4+}$, y25-28$^{4+}$, y30$^{4+}$, y31$^{4+}$, y33-36$^{4+}$ ions were 283.39, 359.18, 384.44, 409.70, 460.22, 477.98, 495.74, 549.02, 584.54, 602.30, 620.06, 637.82, 673.34, 691.10, 726.61, 744.37, 762.13, 779.89 Da. Observed y11$^{5+}$, y16$^{5+}$, y18$^{5+}$, y20$^{5+}$, y22-25$^{5+}$, y27-31$^{5+}$, y33$^{5+}$, y36$^{5+}$ ions were 226.91, 327.96, 368.38, 396.80, 425.21, 439.42, 453.63, 467.83, 496.25, 510.46, 524.66, 538.08, 581.49, 624.11 Da.

In the MS/MS spectrum of 975.70715 Da ion (MH$^{+}$ = 779.80678 Da; putative ID: NANPPAEVVPPQAPAKKAAAAAAAAAAAAAASSSSSSSSCEE), mass/charge ratios of observed b6-9, b13, b14 ions were 565.28, 694.32, 793.38, 892.45, 1285.65, 1382.71 Da, respectively, those of observed b6$^{2+}$, b7$^{2+}$, b9-14$^{2+}$, b17-22$^{2+}$ ions were 283.14, 347.66, 446.73, 495.26, 543.78, 607.81, 643.33, 691.86, 855.47, 890.99, 926.51, 962.03, 997.54, 1033.06 Da. Observed b10$^{3+}$, b14$^{3+}$, b18-20$^{3+}$, b22$^{3+}$, b24$^{3+}$, b25$^{3+}$ ions were 330.51, 461.57, 594.33, 618.01, 641.69, 689.04, 736.40, 760.08 Da. Observed b17-20$^{4+}$, b21$^{4+}$, b22$^{4+}$, b24$^{4+}$, b26$^{4+}$, b28$^{4+}$ ions were 428.24, 446.00, 463.76, 499.26, 517.04, 552.55, 500.07, 605.83 Da. Mass/charge ratios of observed y15-17 ions were 1502.59, 1431.55, 1360.52 Da, respectively, those observed y16$^{2+}$, y18-26$^{2+}$, y32$^{2+}$, y34$^{2+}$ ions were 716.28, 787.32, 822.84, 858.35, 893.87, 929.39, 964.91, 1000.43, 1035.95, 1100.00, 1396.16, 1494.22 Da. Observed y15$^{3+}$, y16$^{3+}$, y22-26$^{3+}$, y28-39$^{3+}$, y41$^{3+}$ ions were 454.18, 477.86, 619.93, 643.61, 667.29, 690.97, 733.67, 800.04, 832.39, 856.07, 898.76, 931.11, 963.46, 996.48, 1029.51, 1072.52, 1096.20, 1128.55, 1160.90, 1222.60 Da. Observed y15$^{4+}$, y23$^{4+}$, y25$^{4+}$, y26$^{4+}$, y29-31$^{4+}$, y35$^{4+}$, y36$^{4+}$, y38$^{4+}$, y40$^{4+}$, y41$^{4+}$ ions were 340.88, 482.96, 518.48, 550.50, 624.55, 642.31, 674.32, 772.38, 804.64, 846.67, 899.44, 917.20 Da.

**Figure S4. Further Evidence Supporting the Role of eRF1 in Regulating C-I30 CTE**

(A) Immunoblots showing the effects of Parkin on ABCE1/Pelo and ABCE1/eRF1 interaction in HeLa cells as shown by co-IP assays.

(B) Immunoblots showing the effect of *PINK1* mutation or OE of NOT4, hMfn1, and VCP-DN on full-length eRF1 level and the generation of a smaller-sized eRF1 fragment (marked with blue arrowhead).

(C) Rescue of the wing posture defect by *eRF1 OE* in *Mhc-Gal4>hMfn1 OE* background. * or #, $p<0.05$ in two-tailed Student's *t*-test vs. *hMfn1-OE* at day 7 or day 15, respectively.

(D) Immunoblots of muscle samples from *PINK1* mutant flies with overexpression of co-translational QC factor (*ABCE1*, *eRF1* and *UPF3*), or with overexpression of co-translational QC factors (*ABCE1*, *eRF1* and *UPF3*) and simultaneous *CI-30 RNAi*, showing the knockdown of C-I30 protein level. Actin serves as loading control.

(E) Wing posture assays of the genotypes shown in D. Wing posture rescue by co-translational QC factor OE was blocked when *CI-30* was knocked down by RNAi. * $p<0.05$ in SNK-test plus Bonferroni correction vs. *control (w-)* group or quality control factor OE at day 7 or day 15, respectively. Note that C-I30 RNAi alone had no effect on wing posture and did not further modify *PINK1* mutant wing posture phenotype.

(F) Immunoblots showing the distribution of mito-dsRed in the total, mito, and cyto fractions. Complex-IV subunit 1 (C-IV s.1) and GAPDH are mito and cyto markers, respectively. KDEL is an ER marker used to show lack of ER contamination in the mito fraction.

(G) Immunoblots showing the distribution of HA-C-I30-Flag in the total, mito, and cyto fractions. Bracket marks the HA-positive but Flag-negative smaller species representing partially synthesized C-I30. Diagram on the right depicts the various products containing the HA and Flag epitopes predicted from the co-translational import model (top) or the cytosolic full-translation followed by mitochondrial import model (bottom).

**Figure S5. Further Characterization of C-I30-CAT-Tail constructs, Effect of Parkin on C-I30-u Aggregate Formation, and Effect of C-I30-u on Proteostasis Signaling**

(A) Immunoblots of C-I30-Flag by 2D gel showing C-I30-Flag-u assembly into the RCC complex in purified mitochondria from HeLa cells transfected with C-I30-Flag. Immunoblots on the top shows C-I30-Flag-u induction by CCCP in the cells used for 2D gel analysis.

(B) Immunoblots of HeLa cell transfected with C-I30-$AT_5$, C-I30-$AT_{23}$ and C-I30-non$AT_{25}$, showing the changes of C-I30-u and the high molecular weight aggregations seen with C-I30-$AT_{23}$ in the absence of CCCP treatment. ] in blue indicates the high MW aggregated species. C-I30-non$AT_{25}$ indicates a C-I30 form with similar lengthened tail as C-I30-$AT_{23}$ but no AT repeats.

(C, D) Effect of transfection of C-I30-no tag, C-I30-3xFlag, or C-I30-$AT_{23}$ on C-I activity (C), and ATP production (D) in HeLa cells. Purified mitochondria are used in these assays. * $p<0.05$, ** $p<0.01$ in SNK-test plus Bonferroni correction vs. vehicle.

(E) Control experiment showing CCCP does not induce aggregation of a GFP-Flag protein in HeLa cells.

(F) Immunostaining of HeLa cells with or without CCCP treatment showing aggregation of endogenous CI-30 by mitochondrial stress. Scale bar, 3μm. Arrowheads: white, aggregates outside mitochondria; yellow: aggregates inside mitochondria.

(G) Immunostaining of HeLa/GFP-Parkin cell transfected with C-I30-Flag and with or without CCCP treatment showing that no aggregate is formed in CCCP treated cells. Scale bar, 3μm.

(H) Qualification of data shown in G, with % of cells with C-I30-Flag aggregates in the total transfected cells indicated. *, $p<0.05$; *Chi-squared* test, compared to non-treated group. #, $p<0.05$; *Chi-squared* test, compared to the transfected HeLa with CCCP treatment. More than 100 transfected cells (FLAG positive) in each group were counted.

(I) Immunoblots showing aggregation of WT C-I30-Flag specifically in CCCP treated cells and WT C-I30-Flag aggregates are enriched in the detergent insoluble fraction.

(J) Immunoblots showing increased p-eIF2α level and reduced p-YAP level in CCCP treated C-I30-Flag transfected cells compared to control cells. p-eIF2α and p-YAP levels are also changed in C-I30-Flag transfected cells without CCCP treatment, consistent with the presence of low level C-I30-Flag-u. Blue arrowhead on C-I30 blot marks endogenous C-I30.

**Figure S6. Further analysis of ARS and Anisomycin Effects on C-I30 Aggregation in Fly Muscle Tissue**

(A) Immunostaining of fly muscle tissue showing that mitochondrial morphology and C-I30 aggregation are not affected by the indicated genetic manipulations in *PINK1* RNAi animals. Scale bar, 5μm. Arrowheads: CI-30 aggregates.

(B, C) Immunostaining of fly muscle tissue showing removal of C-I30 aggregates by Anisomycin treatment in *Mhc-Gal4>hMfn1 OE*, *Mhc-Gal4>VCP DN,* and *Mhc-Gal4>UPF3 RNAi* flies. Arrowheads: CI-30 aggregates.

**Figure S7. Further Characterization of the Mechanism of C-I30-u Formation and Aggregation in Human Cells.**

(A) Immunostaining showing endogenous ATP5a aggregation in CCCP treated HeLa cells (left panels). Control panels on the right show a completely different behavior of SDHA, which accumulates in the cytosol of CCCP treated cells without forming aggregates, consistent with blocked import by CCCP of the cytosolically translated SDHA preprotein (see also Figure 7A).

(B) Immunoblots showing ATP5a-u formation in *PINK1* fly muscle.

(C) Immunostaining showing ATP5a aggregation in *PINK1* fly muscle.

(D) Quantification of data shown in Figure 7B. *, $p<0.05$ in Student's $t$-test compared to Anisomycin non-treatment group.

(E) Immunoblots showing effects of ABCE1 RNAi or OE on C-I30-Flag-u formation in HeLa cells.

(F) Quantification of data shown in Figure 7E. *, $p<0.05$ in Student's $t$-test compared to CCCP non-treatment group in siCon. #, $p<0.05$ in Student's $t$-test compared to siCon after CCCP treatment.

(G) Immunoblots showing effects of the various genetic manipulations of RQC factors on C-I30-Flag-$AT_{23}$ CTE formation and aggregation. Blue ] indicates the aggregated high MW form.

(H) Immunostaining showing effects of the various genetic manipulations on C-I30-Flag-$AT_{23}$ aggregation.

(I) Immunoblots showing effect of Parkin on the recruitment of ANKZF1, NEMF, and eRF1 to MOM-associated 60S ribosomes. Flag (C-I30) blot of HeLa cell samples shows specific accumulation of C-I30-Flag-u on MOM-associated ribosomes, supporting C-I30-Flag-u being an RQC product.

(J) Immunoblots showing accumulation of endogenous C-I30-u and ATP5a-u in *PINK1(G309D)* and *Parkin(C212/V65)* patient fibroblasts compared to fibroblasts from matched control subjects.

(K) Diagram depicting the sequence of events from co-translational quality control of MOM-associated *nRCC* mRNA in response to mitochondrial damage, mRNP remodeling and C-I30 CTE, to C-I30-u aggregate formation, which will seed the formation of disease-defining protein aggregates by sequestering cellular metastable proteins.

**Table S1. Identified Peptides Matching the C-terminus of Fly CI-30 from LC/MS Data**

Peptides highlighted in yellow indicate the peptides containing additional Ala (A) added to the normal C-terminus of CI-30 amino acid sequence. "A" or "A.S" in Red indicate the additional amino acids after the AKK sequence. Observed (M+H) means the observed molecular weight (MW) of the peptides. Mass error (ppm) indicates the difference between the observed MW and theoretical MW.

**Table S2. Putatively Identified Peptides from Customized Pool-Based Searches**

Shown are details of the peptides listed in **Fig. 3I**. Sequences in black indicate the peptides matching the normal peptide sequence of C-I30 protein. Sequences in blue indicate the peptides matching the sequences from the customized database and not from the C-I30 protein sequence. The spectra of selected peptides could be found in **Fig. S3I**. Observed (M+H) means the observed molecular weight (MW) of the peptides. Mass error (ppm) indicates the difference between observed MW and predicted theoretical MW.

Figure S1

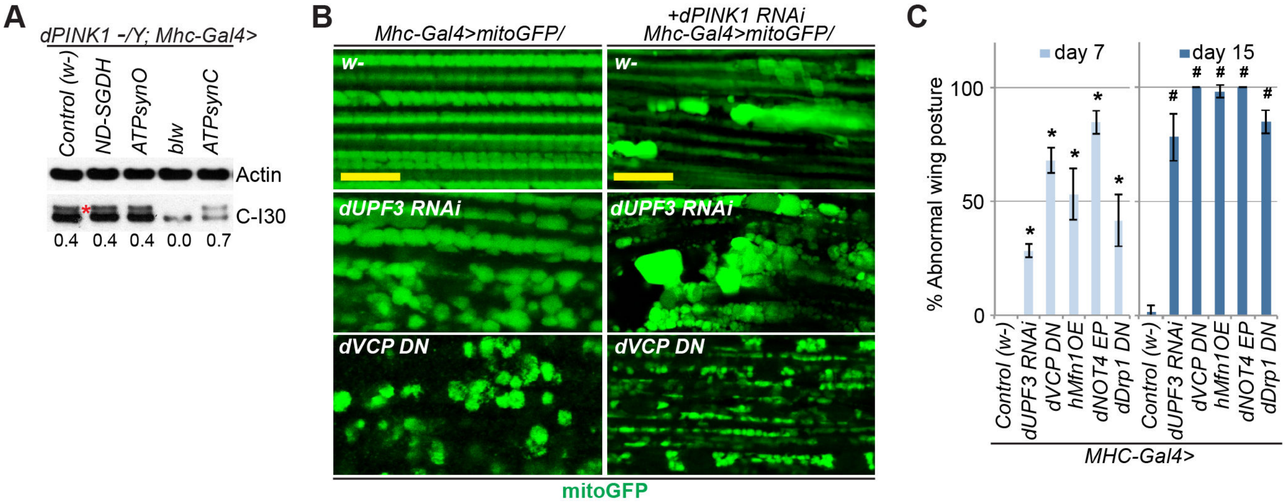
A
dPINK1 -/Y; Mhc-Gal4>
Control (w-)
ND-SGDH
ATPsynO
blw
ATPsynC
Actin
C-I30
0.4
0.4
0.4
0.0
0.7
B
Mhc-Gal4>mitoGFP/
+dPINK1 RNAi
Mhc-Gal4>mitoGFP/
w-
w-
dUPF3 RNAi
dUPF3 RNAi
dVCP DN
dVCP DN
mitoGFP
C
day 7
day 15
% Abnormal wing posture
100
50
0
Control (w-)
dUPF3 RNAi
dVCP DN
hMfn1OE
dNOT4 EP
dDrp1 DN
MHC-Gal4>

Figure S2

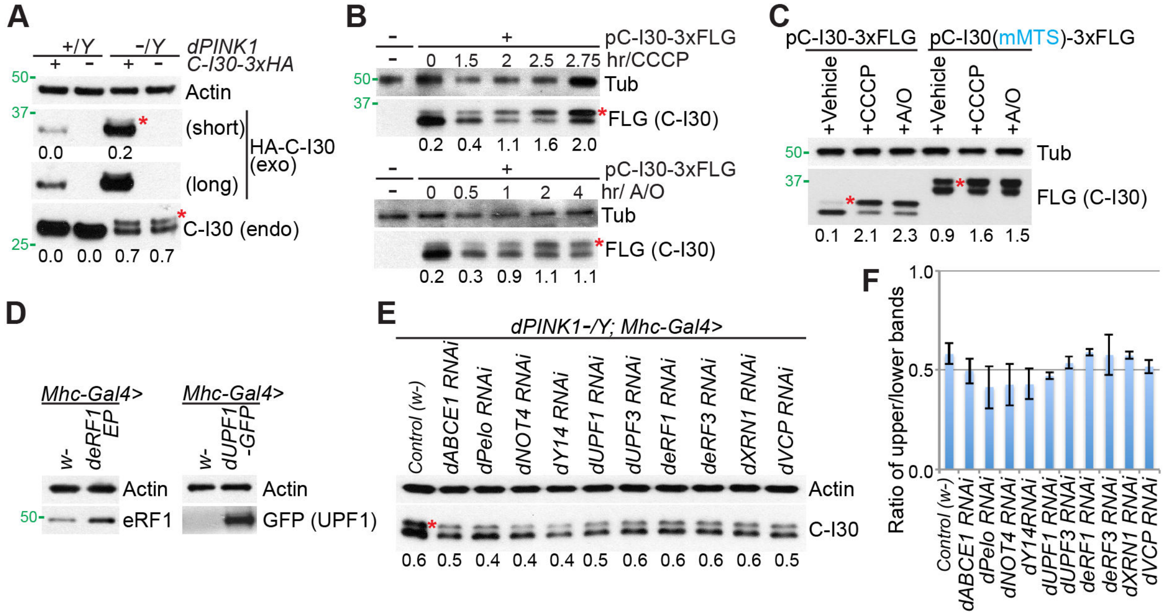
A
+/Y
-/Y
dPINK1
C-I30-3xHA
Actin
(short)
(long)
HA-C-I30
(exo)
C-I30 (endo)
0.0 0.2
0.0 0.0 0.7 0.7
B
pC-I30-3xFLG
hr/CCCP
0 1.5 2 2.5 2.75
Tub
FLG (C-I30)
0.2 0.4 1.1 1.6 2.0
pC-I30-3xFLG
hr/ A/O
0 0.5 1 2 4
Tub
FLG (C-I30)
0.2 0.3 0.9 1.1 1.1
C
pC-I30-3xFLG
pC-I30(mMTS)-3xFLG
+Vehicle
+CCCP
+A/O
Tub
FLG (C-I30)
0.1 2.1 2.3 0.9 1.6 1.5
D
Mhc-Gal4>
w-
deRF1 EP
Actin
eRF1
dUPF1 -GFP
GFP (UPF1)
E
dPINK1-/Y; Mhc-Gal4>
Control (w-)
dABCE1 RNAi
dPelo RNAi
dNOT4 RNAi
dY14 RNAi
dUPF1 RNAi
dUPF3 RNAi
deRF1 RNAi
deRF3 RNAi
dXRN1 RNAi
dVCP RNAi
Actin
C-I30
0.6 0.5 0.4 0.4 0.4 0.5 0.6 0.6 0.6 0.6 0.5
F
Ratio of upper/lower bands
1.0
0.5
0.0
Control (w-)
dABCE1 RNAi
dPelo RNAi
dNOT4 RNAi
dY14RNAi
dUPF1 RNAi
dUPF3 RNAi
deRF1 RNAi
deRF3 RNAi
dXRN1 RNAi
dVCP RNAi

Figure S3

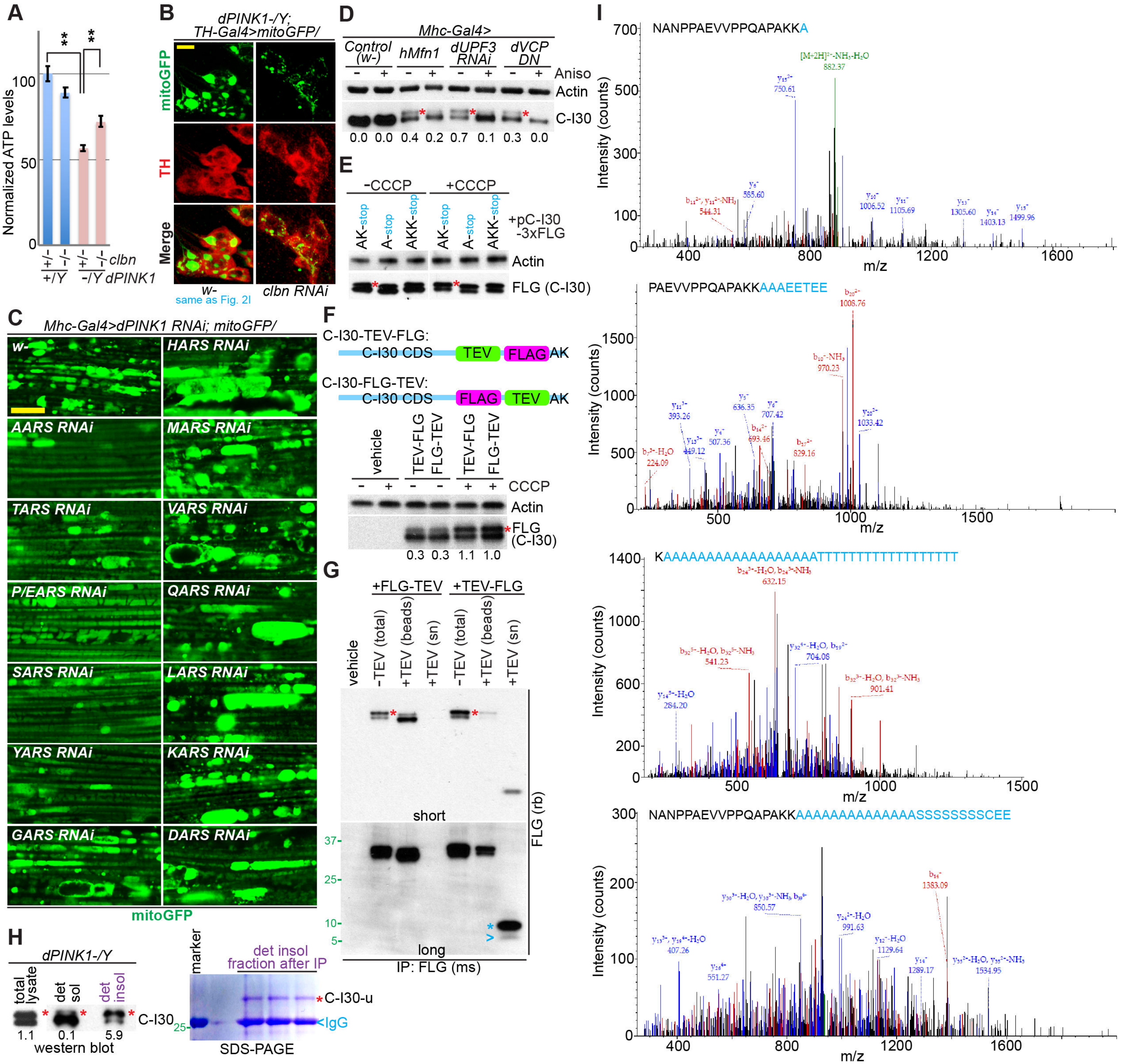

A
Normalized ATP levels
clbn
dPINK1
+/Y
-/Y
B
dPINK1-/Y; TH-Gal4>mitoGFP/
mitoGFP
TH
Merge
w-
same as Fig. 2I
clbn RNAi
C
Mhc-Gal4>dPINK1 RNAi; mitoGFP/
w-
HARS RNAi
AARS RNAi
MARS RNAi
TARS RNAi
VARS RNAi
P/EARS RNAi
QARS RNAi
SARS RNAi
LARS RNAi
YARS RNAi
KARS RNAi
GARS RNAi
DARS RNAi
mitoGFP
D
Mhc-Gal4>
Control (w-)
hMfn1
dUPF3 RNAi
dVCP DN
Aniso
Actin
C-I30
0.0 0.0 0.4 0.2 0.7 0.1 0.3 0.0
E
-CCCP
+CCCP
AK-stop
A-stop
AKK-stop
+pC-I30 -3xFLG
Actin
FLG (C-I30)
F
C-I30-TEV-FLG:
C-I30 CDS
TEV
FLAG
AK
C-I30-FLG-TEV:
vehicle
TEV-FLG
FLG-TEV
CCCP
FLG (C-I30)
0.3 0.3 1.1 1.0
G
+FLG-TEV
+TEV-FLG
-TEV (total)
+TEV (beads)
+TEV (sn)
short
long
FLG (rb)
IP: FLG (ms)
H
dPINK1-/Y
total lysate
det sol
det insol
C-I30
1.1 0.1 5.9
western blot
marker
det insol fraction after IP
C-I30-u
IgG
SDS-PAGE
I
NANPPAEVVPPQAPAKKA
Intensity (counts)
m/z
PAEVVPPQAPAKKAAAEETEE
KAAAAAAAAAAAAAAAAATTTTTTTTTTTTTTTTT
NANPPAEVVPPQAPAKKAAAAAAAAAAAAAASSSSSSSSCEE

Figure S4

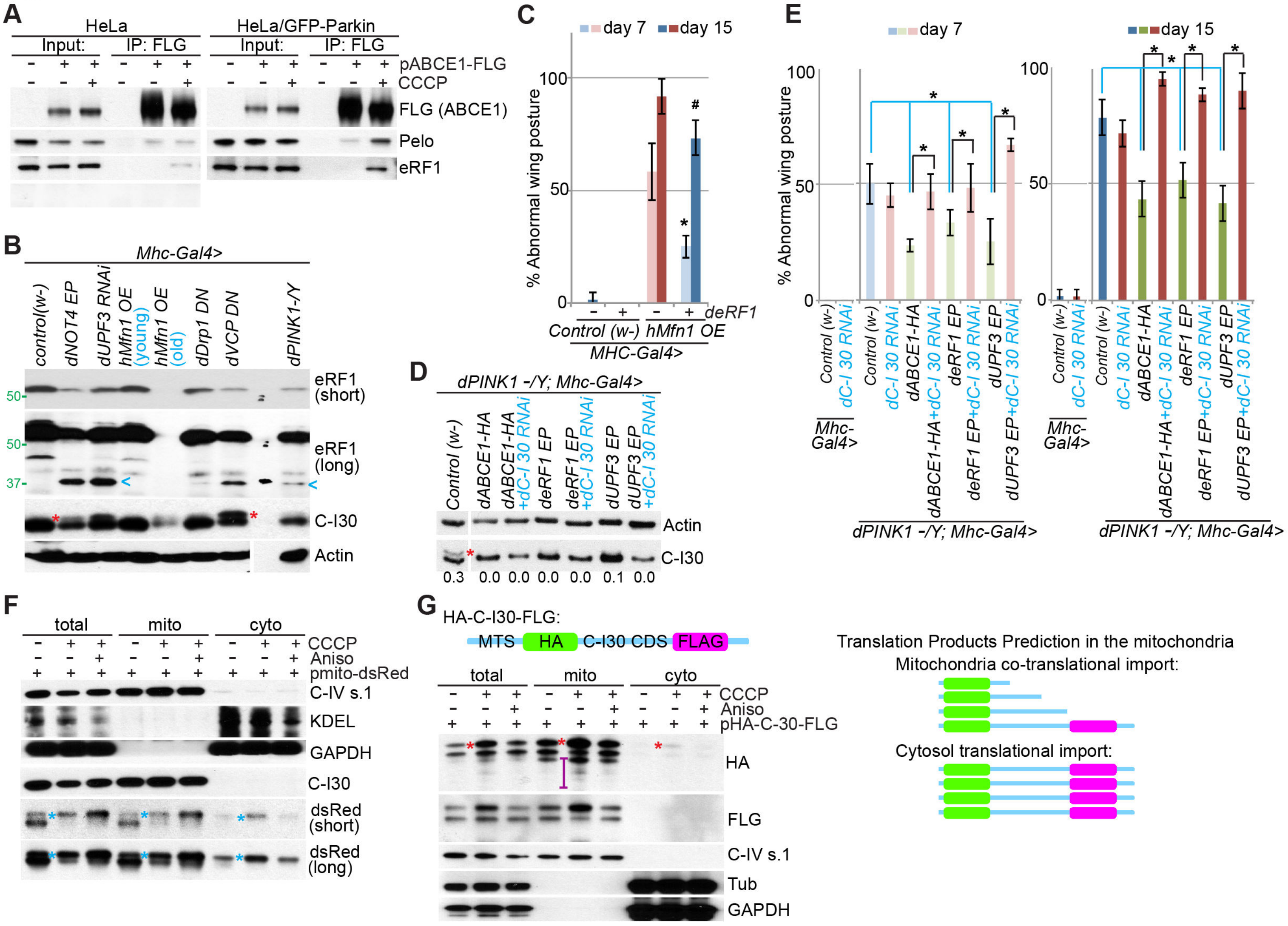
A
HeLa
HeLa/GFP-Parkin
Input:
IP: FLG
pABCE1-FLG
CCCP
FLG (ABCE1)
Pelo
eRF1
B
Mhc-Gal4>
control(w-)
dNOT4 EP
dUPF3 RNAi
hMfn1 OE (young)
hMfn1 OE (old)
dDrp1 DN
dVCP DN
dPINK1-/Y
eRF1 (short)
eRF1 (long)
C-I30
Actin
C
day 7
day 15
% Abnormal wing posture
deRF1
Control (w-)
hMfn1 OE
MHC-Gal4>
D
dPINK1 -/Y; Mhc-Gal4>
Control (w-)
dABCE1-HA
dABCE1-HA +dC-I 30 RNAi
deRF1 EP
deRF1 EP +dC-I 30 RNAi
dUPF3 EP
dUPF3 EP +dC-I 30 RNAi
Actin
C-I30
0.3 0.0 0.0 0.0 0.0 0.1 0.0
E
day 7
day 15
% Abnormal wing posture
Control (w-)
dC-I 30 RNAi
Mhc-Gal4>
dABCE1-HA
dABCE1-HA+dC-I 30 RNAi
deRF1 EP
deRF1 EP+dC-I 30 RNAi
dUPF3 EP
dUPF3 EP+dC-I 30 RNAi
dPINK1 -/Y; Mhc-Gal4>
F
total
mito
cyto
CCCP
Aniso
pmito-dsRed
C-IV s.1
KDEL
GAPDH
C-I30
dsRed (short)
dsRed (long)
G
HA-C-I30-FLG:
MTS
HA
C-I30 CDS
FLAG
total
mito
cyto
CCCP
Aniso
pHA-C-30-FLG
HA
FLG
C-IV s.1
Tub
GAPDH
Translation Products Prediction in the mitochondria
Mitochondria co-translational import:
Cytosol translational import:

Figure S5

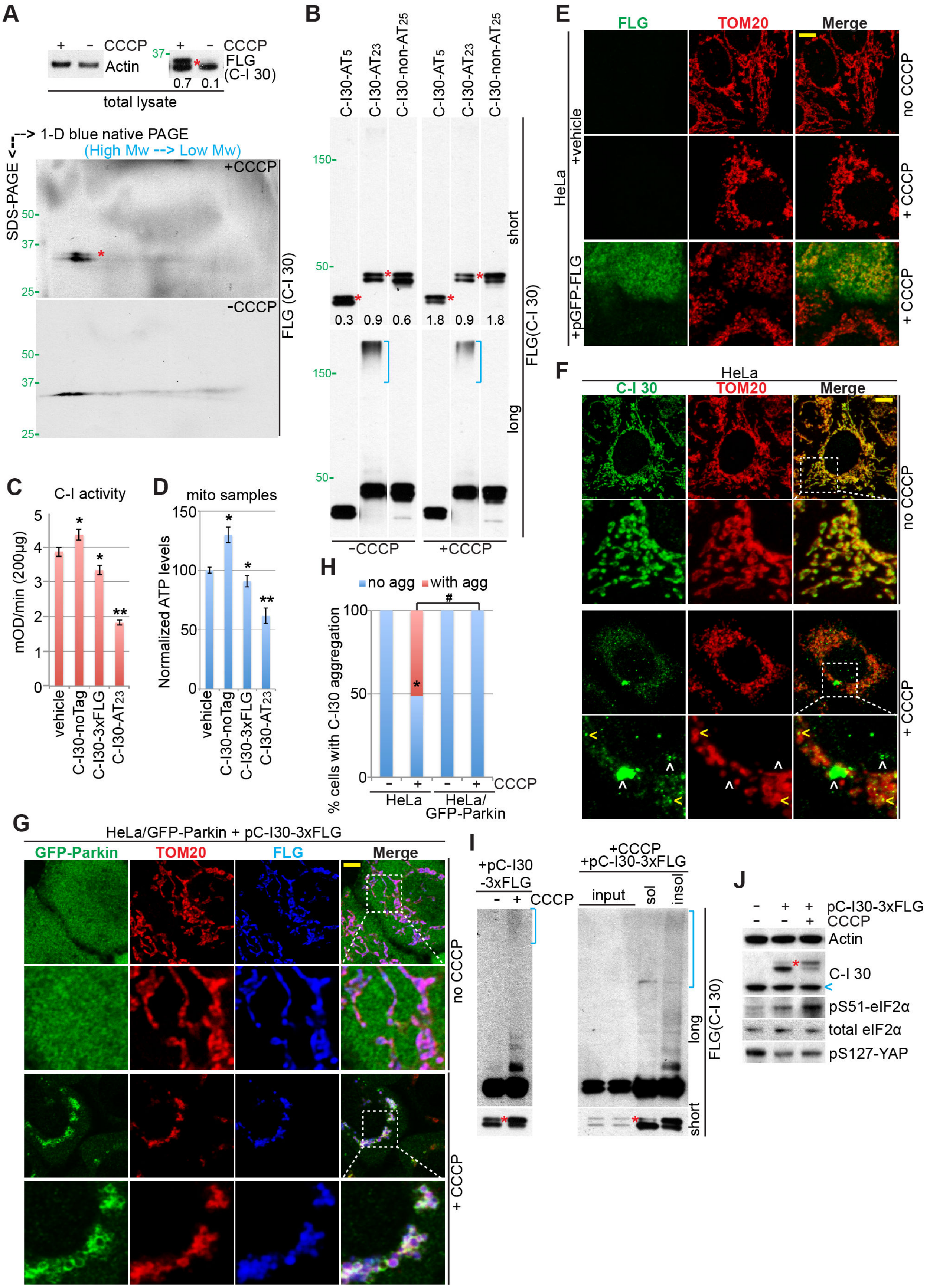
A
+ - CCCP
Actin
37- + - CCCP
FLG (C-I 30)
0.7 0.1
total lysate
1-D blue native PAGE
(High Mw --> Low Mw)
SDS-PAGE
+CCCP
-CCCP
FLG (C-I 30)
B
C-I30-AT5
C-I30-AT23
C-I30-non-AT25
0.3 0.9 0.6 1.8 0.9 1.8
short
long
FLG(C-I 30)
-CCCP
+CCCP
C
C-I activity
mOD/min (200μg)
vehicle
C-I30-noTag
C-I30-3xFLG
C-I30-AT23
D
mito samples
Normalized ATP levels
E
FLG
TOM20
Merge
HeLa
+vehicle
+pGFP-FLG
no CCCP
+ CCCP
F
HeLa
C-I 30
TOM20
Merge
no CCCP
+ CCCP
H
no agg
with agg
% cells with C-I30 aggregation
CCCP
HeLa
HeLa/ GFP-Parkin
G
HeLa/GFP-Parkin + pC-I30-3xFLG
GFP-Parkin
TOM20
FLG
Merge
no CCCP
+ CCCP
I
+pC-I30 -3xFLG
CCCP
+CCCP +pC-I30-3xFLG
input
sol
insol
long
short
FLG(C-I 30)
J
pC-I30-3xFLG
CCCP
Actin
C-I 30
pS51-eIF2α
total eIF2α
pS127-YAP

Figure S6

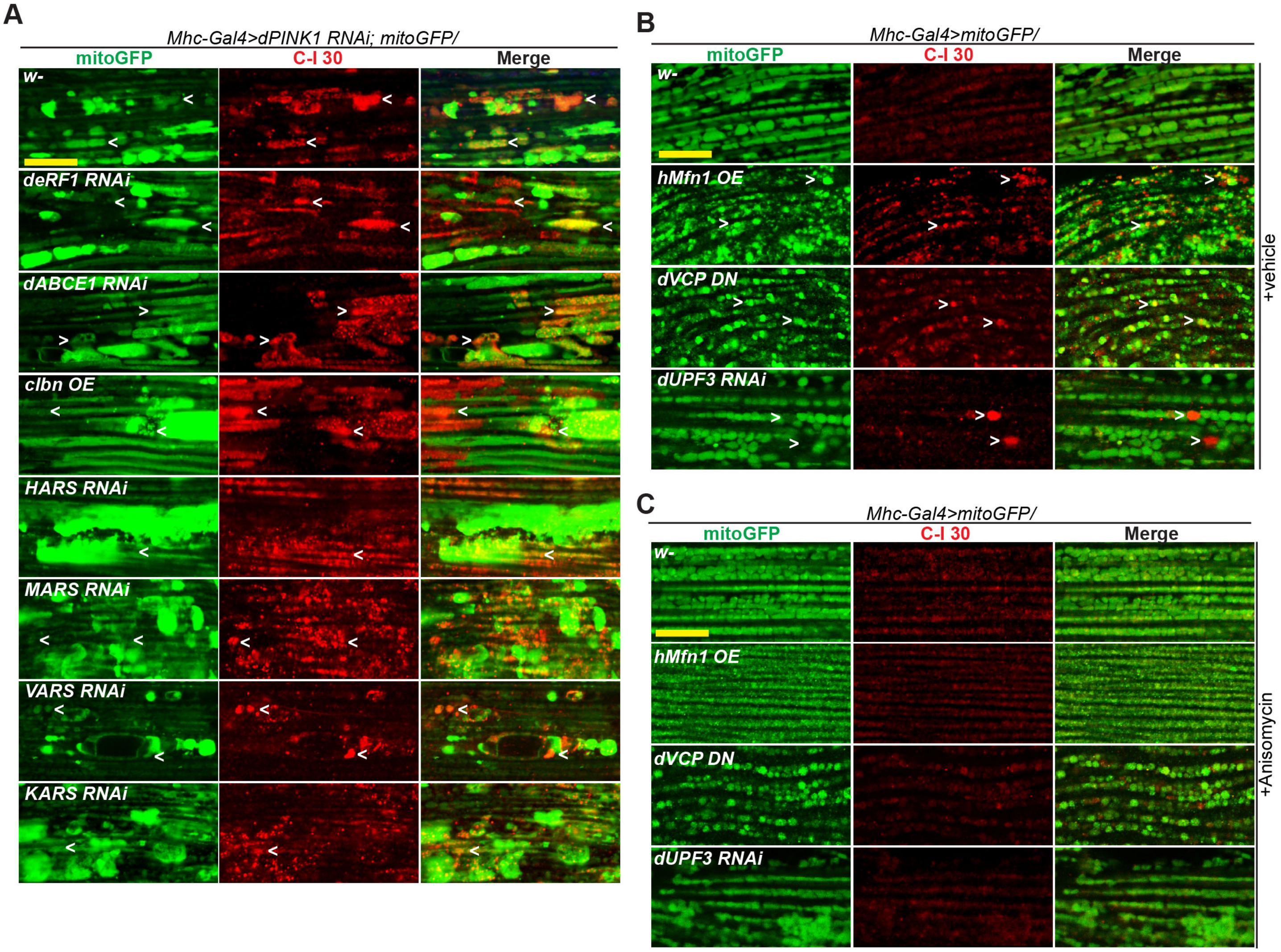
A
Mhc-Gal4>dPINK1 RNAi; mitoGFP/
mitoGFP
C-I 30
Merge
w-
deRF1 RNAi
dABCE1 RNAi
clbn OE
HARS RNAi
MARS RNAi
VARS RNAi
KARS RNAi
B
Mhc-Gal4>mitoGFP/
mitoGFP
C-I 30
Merge
w-
hMfn1 OE
dVCP DN
dUPF3 RNAi
+vehicle
C
Mhc-Gal4>mitoGFP/
mitoGFP
C-I 30
Merge
w-
hMfn1 OE
dVCP DN
dUPF3 RNAi
+Anisomycin

Figure S7

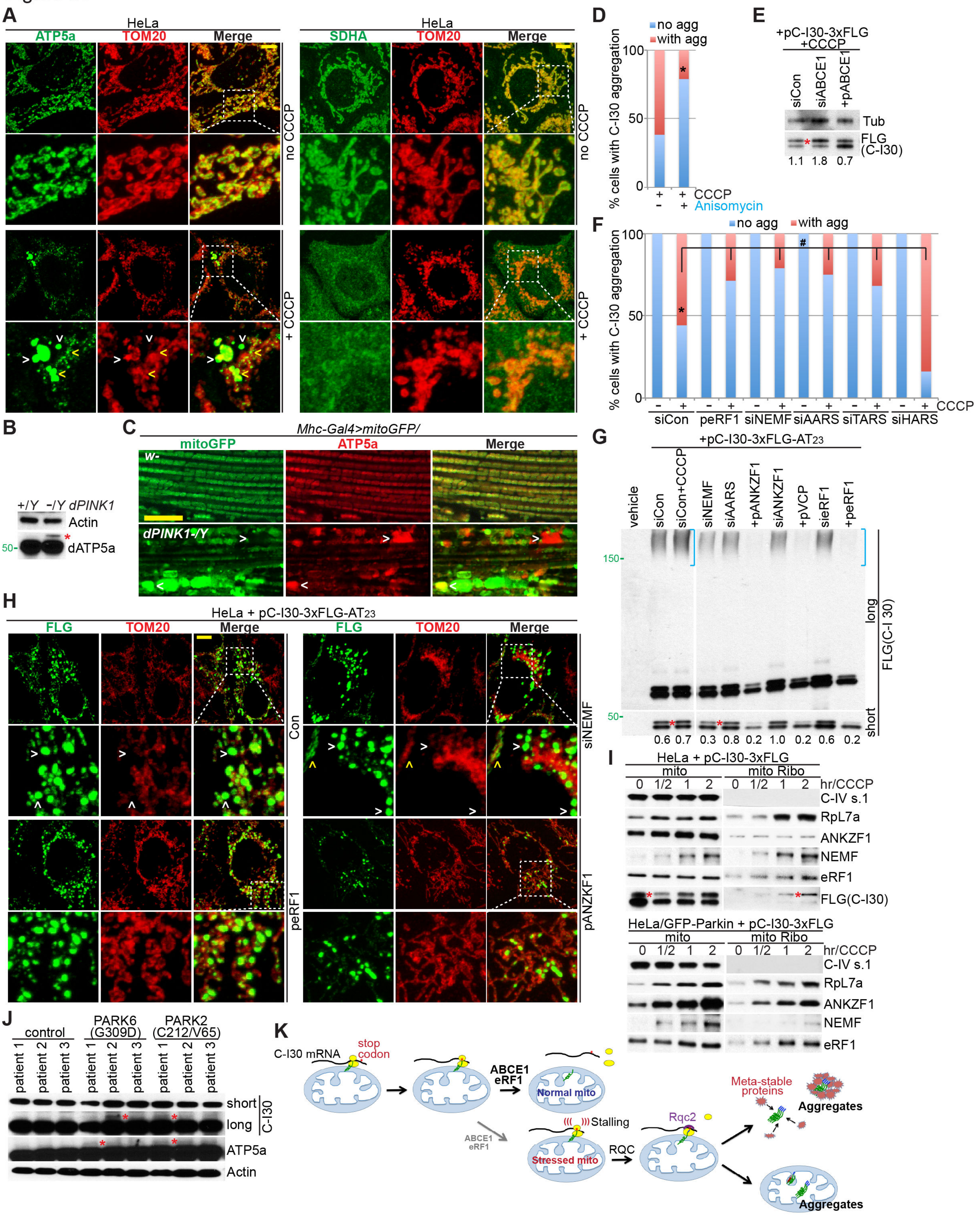

A
HeLa
ATP5a
TOM20
Merge
SDHA
no CCCP
+ CCCP
B
+/Y -/Y dPINK1
Actin
dATP5a
50
C
Mhc-Gal4>mitoGFP/
mitoGFP
ATP5a
Merge
w-
dPINK1-/Y
D
no agg
with agg
% cells with C-I30 aggregation
CCCP
Anisomycin
E
+pC-I30-3xFLG
+CCCP
siCon
siABCE1
+pABCE1
Tub
FLG (C-I30)
1.1 1.8 0.7
F
no agg
with agg
% cells with C-I30 aggregation
CCCP
siCon
peRF1
siNEMF
siAARS
siTARS
siHARS
G
+pC-I30-3xFLG-AT23
vehicle
siCon
siCon+CCCP
siNEMF
siAARS
+pANKZF1
siANKZF1
+pVCP
sieRF1
+peRF1
150
50
long
short
FLG(C-I 30)
0.6 0.7 0.3 0.8 0.2 1.0 0.2 0.6 0.2
H
HeLa + pC-I30-3xFLG-AT23
FLG
TOM20
Merge
Con
siNEMF
peRF1
pANZKF1
I
HeLa + pC-I30-3xFLG
mito
mito Ribo
0 1/2 1 2
hr/CCCP
C-IV s.1
RpL7a
ANKZF1
NEMF
eRF1
FLG(C-I30)
HeLa/GFP-Parkin + pC-I30-3xFLG
J
control
PARK6 (G309D)
PARK2 (C212/V65)
patient 1
patient 2
patient 3
short
long
C-I30
ATP5a
Actin
K
C-I30 mRNA
stop codon
ABCE1
eRF1
Normal mito
Stalling
Stressed mito
RQC
Rqc2
Meta-stable proteins
Aggregates

### Table S1

| Peptide < ProteinMetrics Confidential > | Modifications | Observed (M+H) | Mass error (ppm) | \|Log Prob\| |
|---|---|---|---|---|
| R.KFDLSAPWEQFPNFRNANPPAEVVPPQAPA.K | | 3334.671 | 2.1 | 2.11 |
| R.KFDLSAPWEQFPN[+0.984]FRNANPPAEVVPPQAPA.K | N[+1] | 3335.649 | 0.3 | 0.53 |
| R.KFDLSAPWEQFPNFRN[+0.984]ANPPAEVVPPQAPA.K | N[+1] | 3335.654 | 1.8 | 2.15 |
| R.KFDLSAPWEQFPN[+0.984]FRNANPPAEVVPPQAPA.K | N[+1] | 3335.658 | 3.0 | 5.94 |
| R.KFDLSAPWEQFPNFRN[+0.984]ANPPAEVVPPQAPA.K | N[+1] | 3335.654 | 1.7 | 6.59 |
| R.KFDLSAPWEQFPNFRN[+0.984]ANPPAEVVPPQAPAKK.A | N[+1] | 3591.844 | 1.6 | 0.54 |
| R.KFDLSAPWEQFPNFRN[+0.984]ANPPAEVVPPQAPAKK.A | N[+1] | 3591.846 | 2.1 | 5.06 |
| R.KFDLSAPWEQFPN[+0.984]FRNANPPAEVVPPQAPAKK.A | N[+1] | 3591.847 | 2.4 | 12.00 |
| K.FDLSAPWEQFPNFRNANPPAEVVPPQAPA.K | | 3206.568 | -0.6 | 0.46 |
| K.FDLSAPWEQFPNFRN[+0.984]ANPPAEVVPPQAPA.K | N[+1] | 3207.560 | 2.1 | 5.23 |
| K.FDLSAPWEQFPNFRNANPPAEVVPPQAPAKK.A | | 3462.748 | -3.1 | 2.88 |
| K.FDLSAPWEQFPNFRN[+0.984]ANPPAEVVPPQAPAKK.A | N[+1] | 3463.750 | 2.0 | 4.65 |
| R.NANPPAEVVPPQAPA.K | | 1471.742 | -8.2 | 0.17 |
| R.NANPPAEVVPPQAPA.K | | 1471.752 | -1.6 | 1.47 |
| R.NANPPAEVVPPQAPA.K | | 1471.757 | 2.0 | 2.10 |
| R.NANPPAEVVPPQAPA.K | | 1471.755 | 0.8 | 2.99 |
| R.NANPPAEVVPPQAPA.K | | 1471.755 | 0.5 | 2.91 |
| R.NANPPAEVVPPQAPA.K | | 1471.754 | -0.2 | 2.91 |
| R.NANPPAEVVPPQAPA.K | | 1471.755 | 0.6 | 4.69 |
| R.NANPPAEVVPPQAPA.K | | 1471.755 | 0.9 | 5.30 |
| R.NANPPAEVVPPQAPA.K | | 1471.755 | 0.6 | 7.90 |
| R.NANPPAEVVPPQAPA.K | | 1471.757 | 1.8 | 8.04 |
| R.NANPPAEVVPPQAPA.K | | 1471.756 | 1.4 | 8.27 |
| R.NANPPAEVVPPQAPA.K | | 1471.756 | 1.3 | 9.36 |
| R.NANPPAEVVPPQAPA.K | | 1471.755 | 0.5 | 8.58 |
| R.N[+0.984]ANPPAEVVPPQAPA.K | N[+1] | 1472.743 | 3.4 | 3.00 |
| R.N[+0.984]ANPPAEVVPPQAPA.K | N[+1] | 1472.739 | 0.9 | 4.68 |
| R.N[+0.984]ANPPAEVVPPQAPA.K | N[+1] | 1472.737 | -0.8 | 5.26 |
| R.N[+0.984]ANPPAEVVPPQAPA.K | N[+1] | 1472.740 | 1.2 | 7.88 |
| R.N[+0.984]ANPPAEVVPPQAPA.K | N[+1] | 1472.741 | 1.8 | 8.09 |
| R.NANPPAEVVPPQAPAK.K | | 1599.849 | 0.0 | 5.22 |
| R.NANPPAEVVPPQAPAK.K | | 1599.851 | 1.3 | 9.53 |
| R.NANPPAEVVPPQAPAK.K | | 1599.851 | 1.3 | 9.69 |
| R.NANPPAEVVPPQAPAK.K | | 1599.850 | 0.8 | 12.98 |
| R.NAN[+0.984]PPAEVVPPQAPAK.K | N[+1] | 1600.835 | 1.2 | 2.94 |
| R.N[+0.984]ANPPAEVVPPQAPAK.K | N[+1] | 1600.850 | 10.5 | 4.41 |
| R.N[+0.984]ANPPAEVVPPQAPAK.K | N[+1] | 1600.836 | 2.2 | 5.26 |
| R.N[+0.984]ANPPAEVVPPQAPAK.K | N[+1] | 1600.833 | 0.1 | 8.87 |
| R.N[+0.984]ANPPAEVVPPQAPAK.K | N[+1] | 1600.835 | 1.2 | 9.49 |
| R.NAN[+0.984]PPAEVVPPQAPAK.K | N[+1] | 1600.832 | -0.4 | 12.36 |
| R.NANPPAEVVPPQAPAKK.A | | 1727.947 | 1.6 | 1.60 |
| R.NANPPAEVVPPQAPAKK.A | | 1727.948 | 2.4 | 2.52 |
| R.NANPPAEVVPPQAPAKK.A | | 1727.942 | -0.9 | 4.77 |
| R.NANPPAEVVPPQAPAKK.A | | 1727.945 | 0.6 | 4.97 |
| R.NANPPAEVVPPQAPAKK.A | | 1727.945 | 0.6 | 6.35 |
| R.NANPPAEVVPPQAPAKK.A | | 1727.947 | 2.1 | 5.82 |
| R.NANPPAEVVPPQAPAKK.A | | 1727.945 | 0.5 | 5.83 |
| R.NANPPAEVVPPQAPAKK.A | | 1727.944 | -0.2 | 5.63 |
| R.NANPPAEVVPPQAPAKK.A | | 1727.946 | 1.2 | 6.51 |
| R.NANPPAEVVPPQAPAKK.A | | 1727.945 | 0.6 | 6.60 |
| R.NANPPAEVVPPQAPAKK.A | | 1727.946 | 1.1 | 6.83 |
| R.NANPPAEVVPPQAPAKK.A | | 1727.943 | -0.3 | 6.76 |
| R.NANPPAEVVPPQAPAKK.A | | 1727.944 | 0.2 | 6.09 |
| R.NANPPAEVVPPQAPAKK.A | | 1727.944 | 0.2 | 8.97 |

| | | | | |
|---|---|---|---|---|
| R.NANPPAEVVPPQAPAKK.A | | 1727.945 | 0.5 | 8.81 |
| R.NANPPAEVVPPQAPAKK.A | | 1727.944 | 0.2 | 8.46 |
| R.NANPPAEVVPPQAPAKK.A | | 1727.947 | 1.7 | 10.57 |
| R.NANPPAEVVPPQAPAKK.A | | 1727.946 | 0.9 | 9.28 |
| R.NANPPAEVVPPQAPAK{+14.968}K.A | K{+15} | 1742.912 | 0.0 | 0.53 |
| R.NANPPAEVVPPQAPAK{+14.966}K.A | K{+15} | 1742.910 | 0.0 | 0.75 |
| R.NANPPAEVVPPQAPAK{+58.004}K.A | K{+58} | 1785.948 | 0.0 | 1.53 |
| R.NAN[+0.984]PPAEVVPPQAPAKK.A | N[+1] | 1728.927 | -0.5 | 5.51 |
| R.N[+0.984]ANPPAEVVPPQAPAKK.A | N[+1] | 1728.927 | -0.3 | 6.09 |
| R.NAN[+0.984]PPAEVVPPQAPAKK.A | N[+1] | 1728.927 | -0.3 | 7.14 |
| R.NAN[+0.984]PPAEVVPPQAPAKK.A | N[+1] | 1728.927 | -0.5 | 7.57 |
| R.N[+0.984]ANPPAEVVPPQAPAKK.A | N[+1] | 1728.928 | 0.3 | 9.11 |
| **R.NANPPAEVVPPQAPAKKA.S** | | 1798.986 | 2.9 | 1.47 |
| N.ANPPAEVVPPQAPAKK.A | | 1613.903 | 1.1 | 5.63 |
| A.NPPAEVVPPQAPAKK.A | | 1542.866 | 1.7 | 1.46 |
| A.NPPAEVVPPQAPAKK.A | | 1542.867 | 2.1 | 6.15 |
| N.PPAEVVPPQAPAK.K | | 1300.727 | 0.9 | 6.49 |
| N.PPAEVVPPQAPAK.K | | 1300.727 | 0.9 | 8.62 |
| N.PPAEVVPPQAPAKK.A | | 1428.822 | 0.9 | 1.70 |
| N.PPAEVVPPQAPAKK.A | | 1428.822 | 1.1 | 2.56 |
| N.PPAEVVPPQAPAKK.A | | 1428.824 | 2.3 | 3.35 |
| N.PPAEVVPPQAPAKK.A | | 1428.823 | 1.3 | 4.89 |
| N.PPAEVVPPQAPAKK.A | | 1428.822 | 0.9 | 7.90 |
| N.PPAEVVPPQAPAKK.A | | 1428.822 | 0.9 | 8.44 |

### Table S2

| Peptide < ProteinMetrics Confidential > | Modifications | Observed (M+H) | Mass error (ppm) | q-Value | Spectrum |
|---|---|---|---|---|---|
| NANPPAEVVPPQAPAKKA | | 1798.9861 | 2.79 | 0 | in Fig. S3I |
| PAEVVPPQAPAKKAAAEETEE | | 2162.1116 | 6.41 | 0.002 | in Fig. S3I |
| PPAEVVPPQAPAKKSETCAYCC | | 2289.0591 | -5.34 | 0.002 | |
| RNAnPPAEVVPPQAPAKKAAAAAAAATTTTTTYYYYYYYYCEEEE | N4(Deamidated) | 4983.2915 | -1.38 | 0 | |
| KAAAAAAAAAAAAAAAAAATTTTTTTTTTTTTTTTTT | | 3244.6257 | -4.17 | 0.002 | in Fig. S3I |
| NANPPAEVVPPQAPAKKAAAAAAAAAAAAAASSSSSSSSCEE | | 3779.8068 | -1.99 | 0.002 | in Fig. S3I |
| nAnPPAEVVPPQAPAKKAAAAAAAAATTTTTTEEEE | N1(Deamidated); N3(Deamidated) | 3491.6905 | -3.45 | 0.007 | |
| RNANPPAEVVPPQAPAKKTTTTTTYYYYYYYYCEEEEAAAAAAAA | | 4982.3033 | -2.21 | 0.008 | |
